# TRPML1 positions lysosomes and regulates actin-membrane linkers in astrocyte processes

**DOI:** 10.64898/2026.06.26.734916

**Authors:** Maeve L. Spivey, Madison L. Fuller, Serena J. Chen, David K. Sidibe, Yingqi Wang, Janardhan Bhattarai, Minghong Ma, Sandra Maday

## Abstract

Lysosomes are critical for neuronal physiology and synaptic function, but their organization and roles within astrocytes, an integral component of the tripartite synapse, remain unknown. Here, we use a neuron-astrocyte coculture system that promotes stellate astrocyte morphology to investigate the trafficking of late endosomes and lysosomes (LELs) in astrocytes. As astrocyte branches mature, degradative activity becomes concentrated in the soma and LELs in branches undergo bidirectional motility that becomes progressively attenuated. We establish that the lysosomal cation channel TRPML1 drives LEL immobilization. TRPML1-mediated arrest involves myosin-Va tethering to the actin cytoskeleton, which may position LELs near peripheral astrocyte processes (PAPs), fine actin-enriched protrusions that can contact synapses. Strikingly, TRPML1 activity modulates the phosphorylation of ezrin, radixin, and moesin, actin-membrane linkers enriched in PAPs. These effects are rapid and transient, suggesting a role for lysosomal TRPML1 in regulating PAP membrane dynamics. Thus, TRPML1 positions LELs proximal to PAPs which may influence PAP structural plasticity and astrocyte-synapse contacts.

## INTRODUCTION

Lysosomes are increasingly recognized as key regulators of synaptic function, coordinating protein degradation, secretion, and signaling at pre- and postsynaptic compartments. The transport of lysosomal organelles is tuned to neuronal activity, with late endosome-lysosomes (LELs) stalling near postsynaptic compartments during bouts of heightened synaptic activity (1, 2). This activity-dependent positioning may enable LELs to locally degrade synaptic proteins (2–4), and to secrete lysosomal cathepsins and synaptic organizing molecules that remodel synaptic structure (5, 6). Despite these insights in neurons, how lysosomes are distributed and function within astrocytes, an essential component of the tripartite synapse, remains largely unexplored.

Astrocytes contact a substantial proportion of synapses across brain regions, including estimates of ∼77% of synapses in the striatum (7), ∼86% in the hippocampus (7), and ∼90-99% in the cortex (8, 9). Astrocytes, like neurons, have complex and polarized morphologies that reflect the diversity of synaptic inputs they integrate. Protoplasmic astrocytes exhibit radial symmetry, composed of 4-10 primary branches (1° processes that radiate from the soma), branchlets (2°, 3° and higher order processes), terminal processes that enwrap the vasculature (i.e., endfeet), and a high density of small protrusions that can contact synapses termed peripheral astrocyte processes (i.e., PAPs) (10). PAPs are highly dynamic structures that extend and retract to modulate synaptic coverage and function (9–20). For example, induction of synaptic long-term potentiation (LTP) has been shown to increase PAP motility within minutes of the stimulus, after which PAPs transition to a stabilized, expanded coverage of potentiated synapses (13, 17, 18). However, mechanisms of PAP structural reorganization, or plasticity, may be region and stimulus-specific, as LTP can also induce PAP retraction to enhance glutamate spillover and NMDAR-signaling (15). PAPs are built on a cytoskeletal scaffold of actin and actin-binding proteins, and are further enriched for glutamate and GABA transporters, adhesion molecules, and a diversity of ionotropic and metabotropic neurotransmitter receptors (11, 12, 21). This molecular composition underlies these context-dependent responses, enabling rapid structural remodeling in response to neuronal activity and elevations in intracellular calcium (12–15, 17, 19, 21–23). Given the synaptic demands and morphological complexity of astrocytes, lysosome trafficking and function must be tightly coordinated across their elaborate arbors.

In line with these demands, organelles positive for the lysosome-associated membrane proteins LAMP1/2 have been observed in the soma, branches, and branchlets *in vivo* by both immunostain of endogenous protein (24–26) and detection of exogenously expressed LAMP1 fluorescent fusion proteins (27, 28). Ultrastructural studies have also reported endosomal organelles in astrocyte branches and branchlets *in vivo* (29). Growing evidence implicates astrocytic lysosomes in gliotransmission (30–33), phagocytosis (24, 27), and plasma membrane repair (34). However, LAMP1 labels a heterogeneous organelle population including non-degradative compartments (35–37). Thus, the exact nature of lysosomal organelles, their trafficking dynamics, and functional diversity within astrocyte branches, remain incompletely understood.

The importance of understanding lysosomal biology in astrocytes is underscored by the prominence of neurological deficits across lysosomal storage disorders (LSDs), a group of inherited metabolic diseases caused by mutations in genes encoding lysosomal proteins (38–40). Mucolipidosis type IV (MLIV) is an LSD with early onset neurodegeneration and neurodevelopmental deficits (41, 42) resulting from loss-of-function mutations in *MCOLN1*, the gene encoding TRPML1, a non-selective cation channel widely recognized as a major lysosomal calcium release channel (43–47). Neurological features of MLIV include psychomotor delay, intellectual disability, progressive vision loss, and white matter abnormalities (41, 42). Early descriptions of MLIV patient samples reported prominent lipid accumulations within lysosomes (48, 49), evident in both neurons and glia (49).

Glial pathology has also been reported in a mouse model of MLIV lacking the *MCOLN1* gene (50). In particular, astrocytes accumulated inclusions and exhibited early reactive gliosis that correlated with synaptic deficits (50–52). Proteomic analysis of the cortex from *MCOLN1-/-* mice revealed that compared to control littermates and among other brain cell-types, astrocytes exhibited the largest change in their bulk proteome (53). Notably, *MCOLN1-/-* astrocytes had pronounced changes in actin and synapse-associated proteins, suggesting alterations to astrocyte morphology and PAP function (53). Consistent with this possibility, astrocyte-specific knockout of TRPML1 in the medial prefrontal cortex is sufficient to induce depressive-like behaviors in mice (26), suggesting that astrocytic TRPML1 may shape synaptic function, and in turn, behavior. Yet, how TRPML1 influences lysosome trafficking and function within astrocytic branches is unknown. In neurons and non-neural cells, TRPML1 influences a broad range of lysosomal functions including transport (54–57), degradation (58–60), and exocytosis (26, 61, 62). However, TRPML1 function, and lysosome biology more broadly, is largely unexplored in the context of astrocytes.

Here, we define how TRPML1 influences lysosome trafficking and function in astrocytes using a neuron-astrocyte coculture system that recapitulates key aspects of astrocyte maturation and neuron-astrocyte interactions *in vivo*. We find that as astrocyte branches mature in coculture with neurons, LELs undergo a spatial reorganization. Degradative activity becomes concentrated in the soma, while LELs within branches exhibit short-range bidirectional motility that is progressively attenuated during maturation. We identify TRPML1 as a key regulator of LEL dynamics in astrocyte branches. Pharmacological activation of TRPML1 reduces LEL motility in astrocyte branches, whereas pharmacological inhibition or loss of TRPML1 increases motility. TRPML1-mediated arrest involves myosin-Va tethering to the actin cytoskeleton, which may position LELs near PAPs. In fact, TRPML1 activity rapidly regulates the phosphorylation state of actin-membrane linkers ezrin-radixin-moesin (i.e., ERM proteins), key structural components of PAPs whose phosphorylation state is linked to membrane dynamics. Together, these findings support a model in which TRPML1 coordinates lysosomal positioning along astrocyte branches, enabling LELs to locally regulate actin-membrane linkers that influence PAP structure and plasticity.

## METHODS

### Reagents

Primary antibodies for immunofluorescence include chicken anti-GFP (1:500 dilution; Aves Lab Inc., GFP-1020), chicken anti-GFAP (1:500 dilution; Aves Lab Inc., GFAP), rabbit anti-GFAP (1:1,000 dilution; Millipore, AB5804), rabbit anti-MAP2 (1:100 dilution; Millipore Sigma, ab5622), mouse anti-β3 tubulin (1:100 dilution; R&D Systems, MAB1195), rabbit anti-EAAT1/GLAST (1:200 dilution; Abcam, ab416), mouse anti-S100β (1:100 dilution; Sigma, S2532), rabbit anti-Aquaporin-4 (1:200 dilution; Abcam, HPA014784), rabbit anti-phospho-Ezrin (Thr567)/Radixin (Thr564)/Moesin (Thr558) (p-ERM; 1:200 dilution; Cell Signaling, 3726), rabbit anti-IBA1 (1:100 dilution; Wako, 019-19741), mouse anti-O4 (1:5 dilution; supernatant generously gifted from Dr. Judy Grinspan), and mouse anti-A2B5 (1:2 dilution; hybridoma from ATCC CRL-1520; clone 105). Primary antibodies for immunoblotting include rabbit anti-pERM (1:500-1,000 dilution; Cell Signaling, 3726), rabbit anti-ERM (1:500-1,000 dilution; Cell Signaling, 3142), rabbit anti-GAPDH (1:4,000-7,000 dilution; Cell Signaling, 2118S), and mouse anti-GAPDH (1:3,000 dilution; Advanced Immunochemicals, 2-RGM-2, clone 6C5).

Secondary antibodies for immunofluorescence were used at a 1:200 dilution and include goat anti-chicken Alexa Fluor 488 (Jackson ImmunoResearch, 103-545-155), goat anti-mouse Alexa Fluor 488 (Invitrogen, A11029), goat anti-rabbit Alexa Fluor 488 (Invitrogen, A11034), goat anti-mouse Rhodamine (TRITC) (Jackson ImmunoResearch, 115-025-075), goat anti-rabbit Alexa Fluor 594 (Invitrogen, A11037), goat anti-mouse Alexa Fluor 594 (Invitrogen, A11032), goat anti-rabbit Alexa Fluor 647 (Invitrogen, A21245), goat anti-mouse Alexa Fluor 647 (Invitrogen, A32728), and goat anti-rat Alexa Fluor 647 (Invitrogen, A21247). Secondary antibodies for immunoblotting included donkey anti-rabbit peroxidase-conjugated (1:2,000 dilution; Jackson ImmunoResearch, 711-035-152), and donkey anti-mouse peroxidase-conjugated (1:2,000 dilution; Jackson ImmunoResearch, 711-035-151).

Reagents used include Hoechst 33342 (Molecular Probes, H3570), SiR-Lysosome (Cytoskeleton, CY-SC012), LysoTracker Red (Invitrogen, L7528), DQ-Red-BSA (Thermo Fisher Scientific/Molecular Probes; D12051), BSA-Alexa Fluor 647 (Thermo Fisher Scientific/Molecular Probes; A34785), Fluo-4 AM (Invitrogen, F14201), ML-SA1 (Tocris, 4746; MedChemExpress, HY-108462), ML-SI3 (MedChemExpress, HY-139426; MedKoo Biosciences, 556041), MK6-83 (MedChemExpress, HY-110238), Latrunculin-A (LatA; Tocris, 3973100U), Pepstatin-A (Sigma-Aldrich, P5318-5MG), Aloxistatin (E64d; MedChemExpress, HY-100229), 6-cyano-7-nitroquinoxaline-2,3-dione (CNQX; Tocris, 0190), and (2R)-amino-5-phosphonovaleric acid (AP5; Tocris, 0106).

Lentivirus expression plasmids designed on VectorBuilder include pLV[Exp]-GFAP(short)>EGFP (VB240314-1622dkx), pLV[Exp]-GFAP(short)>GCaMP6s(ns):3xGGGGS:mMcoln1[NM_053177.1] (VB230428-1224kck), pLV[Exp]-GFAP(short)>mCherry/3xGGGGS/mMcoln1[NM_053177.1] (VB231018-1455xzj), pLV[Exp]-GFAP(short)>pEGFP-MyosinVa-Cterminal (VB231128-1184jtc; sequence for the ORF was obtained from Addgene plasmid #110169) (63), pLV[Exp]-GFAP(short)>LifeAct-mNeonGreen (VB250227-1340yef; sequence for the ORF was obtained from Addgene plasmid #98877) (64), pLV[Exp]-GFAP(short)>mLamp1[NM_001317353.1]/3xGGGGS/mCherry (VB240205-1264wfq), pLV[Exp]-GFAP(short)>EB3-mNeonGreen (VB250613-1454xpz; sequence for the ORF was obtained from Addgene plasmid #98881) (65). Lentivirus miR30-shRNA plasmids designed on VectorBuilder include pLV[miR30]-GFAP(short)>EGFP:Scramble[miR30-shRNA#1] (VB230624-1114szn; Target Sequence: ACCTAAGGTTAAGTCGCCCTCG), pLV[miR30]-GFAP(short)>EGFP:mMcoln1[miR30-shRNA#1] (VB230426-1624ggf; Target Sequence: CGCCGCCTCAAGTACTTCTTTA), and pLV[miR30]-GFAP(short)>EGFP:mMcoln1[miR30-shRNA#2] (VB230426-1625ymx; Target Sequence: AGATGATTGCCACTAGACTTTA). All VectorBuilder constructs were validated by DNA sequencing using Plasmidsaurus. Lentivirus plasmid pMDLg/pRRE was a gift from Didier Trono (Addgene plasmid # 12251; http://n2t.net/addgene:12251 ; RRID:Addgene12251) (66). CMV-VSV-G was a gift from Bob Weinberg (Addgene plasmid # 8454; http://n2t.net/addgene:8454 ; RRID:Addgene8454) (67). pRSV-Rev was a gift from Didier Trono (Addgene plasmid # 12253; http://n2t.net/addgene:12253; RRID:Addgene_12253) (66).

### Mouse Line

Transgenic mice expressing GFP-LC3 (eGFP fused to the C-terminus of rat LC3B) were obtained from the RIKEN BioResource Research Center (RBRC00806; strain name B6.Cg-Tg(CAG-eGFP/LC3)53Nmi/NmiRbrc; GFP-LC3#53) and maintained as heterozygotes. The GFP-LC3 transgene is widely expressed using a constitutive CAG composite promoter that combines a cytomegalovirus immediate early enhancer, a chicken β-actin gene promoter and a rabbit β-globin splice acceptor (68). All animal protocols were approved by the Institutional Animal Care and Use Committee at the University of Pennsylvania.

### Primary Cortical Neuron-Astrocyte Coculture

Primary cortical neurons were isolated from the cerebral cortices of non-transgenic mouse embryos of either sex at embryonic day 15.5, as previously described (1). Cortices were digested with 0.25% trypsin for 10 minutes at 37°C and triturated to generate a homogeneous cell suspension. Neurons were plated at 3 million cells per 10 cm dish filled with 8 x 25 mm acid-washed Deckgläser coverslips, or 1.15 million cells per 6 cm dish filled with 5 x 18mm Deckgläser coverslips coated with 100 µg/mL poly-L-lysine (Sigma, P2636). Neurons were plated in attachment media (MEM [Gibco, 11095-072] supplemented with 10% heat-inactivated horse serum [Gibco, 16050-122], 1 mM sodium pyruvate [Gibco, 11360-070], 33 mM glucose [Sigma, G8769], and 37.5 mM NaCl) for 1-4 hours at 37°C in a 5% CO₂ incubator. Coverslips were then transferred to individual wells of a 6-well plate and maintained for 4-5 DIV in neuron maintenance media (Neurobasal medium [Thermo Fisher Scientific/Gibco, 21-103-049] supplemented with 2% B-27 [Thermo Fisher Scientific/Gibco, 17504-044], 37.5 mM NaCl, 33 mM glucose [Sigma, G8769], 2 mM GlutaMAX [Thermo Fisher Scientific/Gibco, 35050-061], and 100 U/mL penicillin and 100 µg/mL streptomycin [Thermo Fisher Scientific/Gibco, 15140-122]) prior to coculturing with astrocytes.

Primary cortical astrocytes were isolated from the cerebral cortices of GFP-LC3 transgenic or non-transgenic neonatal mice of either sex at postnatal day 0-1, as previously described (1). GFP-LC3 transgenic brain tissue was distinguished from non-transgenic brain tissue at the start of the dissection based on the presence or absence of GFP fluorescence, respectively. Astrocytes were plated at 1.5-3 million cells per 10 cm cell culture dish (Corning® Primaria™, 353803) and maintained in glial medium (DMEM [Thermo Fisher Scientific/Gibco, 11965-084] supplemented with 10% heat-inactivated fetal bovine serum [Hyclone, SH30071.03], 2 mM GlutaMAX [Thermo Fisher Scientific/Gibco, 35050-061], and 100 U/mL penicillin and 100 µg/mL streptomycin [Thermo Fisher Scientific/Gibco, 15140-122]) at 37°C in a 5% CO₂ incubator. Primary astrocytes received a full media replacement with fresh, pre-warmed glial media the following day, and every 3-4 days after. Dishes were vigorously tapped to dislodge microglia before aspirating and discarding glial media during media replacements or passaging cells.

At 70-90% confluency, astrocytes were dissociated from 10 cm tissue culture dishes using 0.05% trypsin-EDTA (Gibco, 15400-054). Glial media was added to inactivate the trypsin and astrocytes were spun down at 644 *x g* for 2 minutes. For monoculture experiments, astrocytes were resuspended in glial media and plated at 120,000-175,000 cells per well of a 6-well plate for immunoblotting, or per 35-mm FluoroDish (World Precision Instruments, FD35-100) coated with 500 µg/mL poly-L-lysine (Sigma, P2636) for live-cell microscopy, and maintained until reaching ∼70-80% confluency. For coculture experiments, astrocytes were resuspended in coculture medium (Neurobasal Medium supplemented with 2% B-27, 1% G-5 [Gibco, 17503012], 0.25% [0.5 mM] GlutaMAX, 100 U/mL penicillin, and 100 µg/mL streptomycin). Once monocultured neurons reached DIV4-5, neuronal maintenance medium was aspirated and astrocytes (resuspended in cocultured medium) were plated onto the neuron culture at a density of ∼40,000-67,500 astrocytes per 6-well. After 1-2 DIV of coculture, 2 µM cytosine β-D-arabinofuranoside (AraC; Sigma, C6645) was added to inhibit astrocyte proliferation. Cocultures were maintained for 3-10 DIV depending on the experiment.

### HEK-293T Culturing and Third-Generation Lentivirus Production

HEK-293T cells (ATCC, CRL-3216) were maintained in HEK media (DMEM supplemented with 10% heat-inactivated fetal bovine serum, 2 mM GlutaMAX, 100 U/mL penicillin, and 100 µg/mL streptomycin), and grown on 10 cm tissue culture dishes at 37°C in a 5% CO₂ incubator, as previously described (69). HEK-293T cells were screened annually for mycoplasma contamination by two services: ATCC Cell Authentication Testing Service (Manassas, VA; RRID:SCR_001672) or the University of Pennsylvania Cell Center Services Core Facility (Philadelphia, PA; RRID:SCR_022391). No mycoplasma contamination was detected throughout the duration of the experiments in this manuscript.

At 70-80% confluency, HEK-293T cells were dissociated from the dish using 0.05% trypsin-EDTA. HEK media was added to inactivate the trypsin, and HEK-293T cells were spun down at 644 *x g* for 2 minutes. Cells were resuspended in HEK media and plated at 150,000-350,000 cells per well in 6-well dishes. At 30-40% confluency, HEK-293T cells were transfected using FuGENE 6 (Promega, F6-1000) according to the manufacturer’s instructions. For lentiviral packaging, HEK-293T cells were co-transfected with pCMV-VSV-G (0.5 µg/well; Addgene #8454), pMDLg/pRRE (0.5 µg/well; Addgene #12251), and pRSV-REV (0.5 µg/well; Addgene #12253), together with either shRNAmiR plasmids (1 µg/well) or plasmids encoding fluorescently tagged proteins of interest (1.5 µg/well). In experiments requiring co-transduction of two lentiviruses, viruses were generated separately and then combined at the time of transduction, rather than pooling viral vectors at the DNA transfection stage.

The day following transfection, HEK-293T cell medium was replaced with pre-warmed coculture medium. After 24 hours, lentivirus-containing supernatants were collected and clarified by centrifugation at 644 *x g* for 15 minutes to pellet cellular debris. Clarified supernatants were applied to cocultures at a 1:10 dilution, 3-5 days prior to imaging to allow sufficient time for transgene expression. Remaining supernatants were stored at -80°C.

All recombinant lentiviruses used are third-generation lentiviruses and are replication-incompetent. Work involving recombinant lentivirus was conducted under biosafety level 2 (BSL-2) conditions in accordance with protocols approved by the Institutional Biosafety Committee (under the Environmental Health and Radiation Safety division) of the University of Pennsylvania.

### Microscopy

Imaging of live-cell and fixed-cell immunofluorescence was performed on a BioVision spinning disk confocal microscope system comprised of a Leica DMi8 inverted widefield microscope, a Yokogawa W1 spinning-disk confocal unit, and a Photometrics Prime 95B sCMOS camera. The system was equipped with an environmental chamber to maintain 37°C for live-cell imaging and adaptive focus control to maintain a constant focal plane during time-lapse acquisition. For live-cell imaging, 18 mm coverslips were mounted in a Chamlide CMB magnetic imaging chamber (BioVision Technologies) and imaged for a maximum of 30-40 minutes in HibE imaging medium (Hibernate E [Brain Bits, HE-Lf] supplemented with 2% B-27 and 2 mM GlutaMAX). All images were acquired using VisiView software. In Figures 1A-D, 2A, 3B-E, 4B, 5A-Fi, S2A-E, S3B-I, S5A-C, S7I-K, we used a 63X/1.4 NA plan apochromat oil-immersion objective. In Figures 1E-N, 2C-I, 3F-I, 4C-J, S2F-G, S4A-H, S5I-K, S6A-D, we used a 63X/1.4 NA plan apochromat oil-immersion objective with a 2X optical magnifier. In Fig. S1B-E and Fig. S5D-Di, we used a 40X/1.3 NA plan apochromat oil-immersion objective. Samples were imaged with solid-state 405, 488, 561, and 640 nm lasers for excitation.

**Figure 1.**
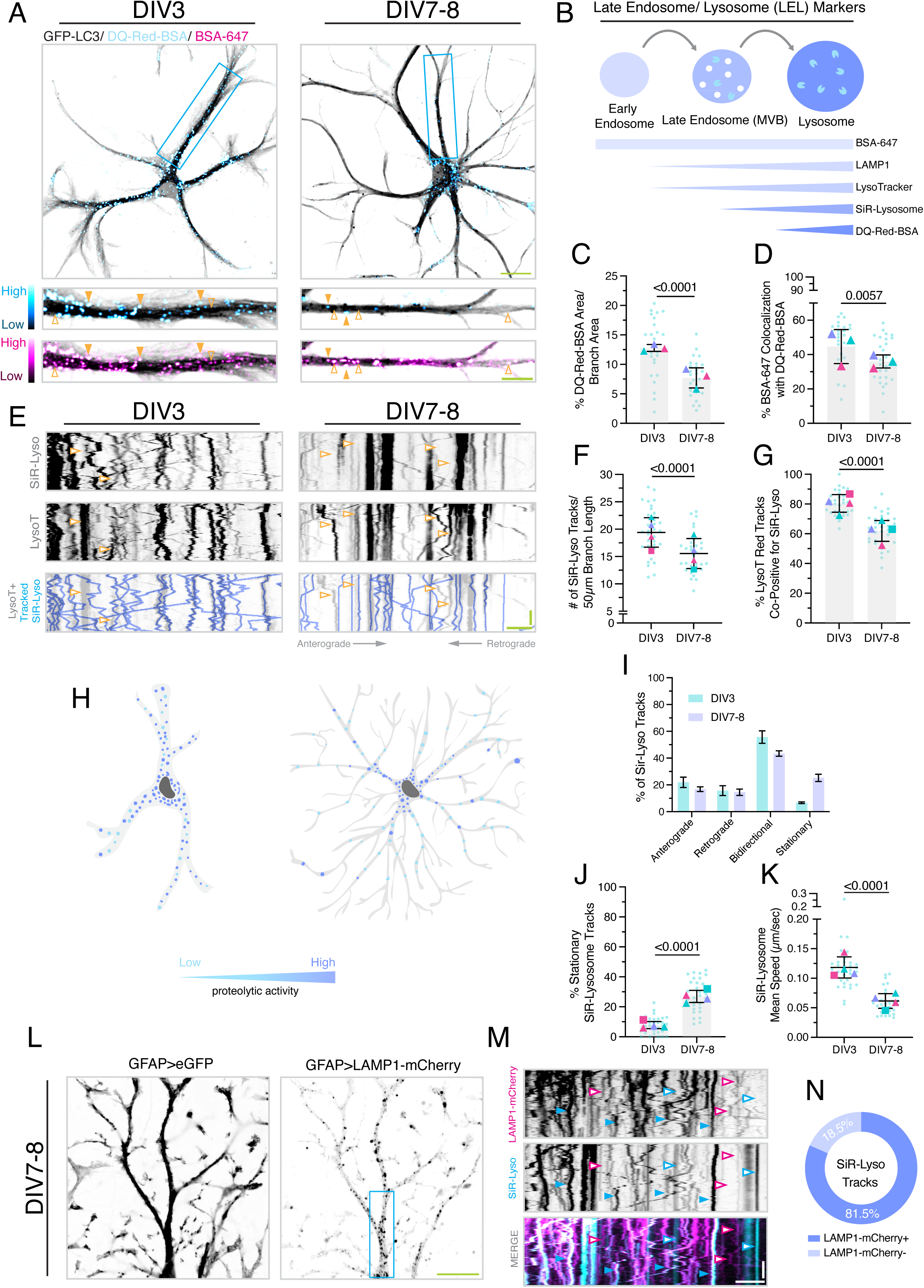
During astrocyte maturation, LELs establish a spatial degradative gradient, with degradative activity concentrating in the soma and branch LEL motility becoming progressively attenuated. (A) Maximum-intensity projections of z-stacks acquired of GFP-LC3 transgenic astrocytes labeled with DQ-Red-BSA and BSA-647 at DIV3 or DIV7-8 of coculture. To visualize LELs in astrocytes, the inverted grayscale GFP-LC3 image was merged with the pseudocolored DQ-Red-BSA or BSA-647 image. Image of entire astrocyte; scale bar, 20 µm. Boxed regions correspond to straightened segments of the astrocyte branch shown below; scale bar, 5 µm. Images within each channel are displayed with identical minimum and maximum intensity settings across DIV. Filled orange arrowheads denote BSA-647 puncta that are co-positive for DQ-Red-BSA. Open orange arrowheads denote BSA-647 puncta that are negative for DQ-Red-BSA. (B) Schematic of probes used in this study to label the spectrum of organelles in the endolysosomal system. (C-D) Corresponding quantification of (A) to measure (C) the total area occupied by DQ-Red-BSA-positive puncta normalized to branch area and (D) the percentage of total BSA-647-positive puncta area that is co-positive for DQ-Red-BSA. Horizontal bars represent the mean of biological replicates ± SD; shown are p-values from a LME model; N=25-26 astrocytes from 3 biological replicates. (E) Representative kymographs of SiR-Lysosome (SiR-Lyso) and LysoTracker Red (LysoT) motility along a primary-to-secondary astrocytic branch from GFP-LC3 transgenic astrocytes in coculture. SiR-Lysosome puncta that were manually tracked (shown in periwinkle) are overlaid onto the LysoTracker Red kymographs. Open orange arrowheads denote LysoTracker Red tracks that are negative for SiR-Lysosome. All quantification of LEL dynamics is derived from kymograph analysis. Horizontal bar, 5 µm; vertical bar, 30 sec. (F-G) Corresponding quantification of (E) to measure (F) SiR-Lysosome track density normalized to a 50 µm branch segment and (G) percentage of LysoTracker Red tracks that are co-positive for SiR-Lysosome. Horizontal bars represent the mean of biological replicates ± SD; shown are p-values from a LME model; N=33-34 branches (one branch per astrocyte) across 4 biological replicates. (H) Schematic depicting the spatial distribution of degradative activity in cocultured astrocytes at DIV3 versus DIV7-8. (I) Corresponding quantification of (E) to measure directionality of SiR-Lysosome motility at DIV3 and DIV7-8 of coculture. Shown are means ± SD; N=20-23 branches (one branch per astrocyte) across 3 biological replicates. (J-K) Corresponding quantification of (E) to measure (J) percentage of stationary (or immobile) tracks and (K) mean speed of SiR-Lysosome tracks. Horizontal bars represent the mean of biological replicates ± SD; shown are p-values from a LME model; N=30-33 branches (one branch per astrocyte) across 4 biological replicates. Data in E-G and I-K are derived from the DMSO control data at each timepoint shown in Figure S4A-D. (L-N) (L) Representative single-plane images of a non-transgenic astrocytic branch transduced with lentiviral eGFP and LAMP1-mCherry (both driven under a shortened GFAP promoter), cocultured for DIV7-8. Scale bar, 20 µm. (M) Corresponding kymographs of LAMP1-mCherry and SiR-Lysosome motility along the astrocyte branch that is boxed in (L). Closed blue arrowheads denote SiR-Lysosome tracks that are positive for LAMP1-mCherry. Open blue arrowheads denote SiR-Lysosome tracks that are negative for LAMP1-mCherry. Open pink arrowheads denote LAMP1-mCherry tracks that are negative for SiR-Lysosome. Horizontal bar, 5 µm; vertical bar, 30 sec. (N) Corresponding quantification of the percentage of SiR-Lysosome tracks that are co-positive for LAMP1-mCherry. N=1,267 tracks from 33 branches (one branch per astrocyte) across 5 biological replicates. Data in L-N are derived from the DMSO control data shown in Figure S4E-H. For all superplots in the paper, small dots indicate the measurements from individual cells (e.g., the technical replicates) from each of the independent experiments; large shapes (e.g. triangles and squares) indicate the corresponding mean from each independent experiment (e.g., the biological replicates).

### Immunostaining

On DIV3-10 of coculture, samples were fixed for 8 min in 4% paraformaldehyde (PFA; Sigma, P6148) and 4% sucrose (Fisher Scientific, BP220-1) in PBS (50 mM NaPO₄ pH 7.4, 150 mM NaCl); fixative was pre-warmed to 37°C. Coverslips were washed twice with PBS and permeabilized for 5 min in 0.1% Triton X-100 (Thermo Fisher Scientific, BP151-100) in PBS, and washed twice with PBS. For experiments in which F-actin was labeled with lentiviral LifeAct-mNeonGreen (Fig. 5A-B), coverslips were permeabilized with ice-cold 100% methanol for 10 minutes, followed by extensive washing with PBS to rehydrate cells.

Following permeabilization, coverslips were blocked for 1 hour in PBS containing 5% goat serum (Sigma, G9023) and 1% BSA (Thermo Fisher Scientific, BP1605-100). Primary antibodies diluted in blocking solution were then added to coverslips and incubated for 1 hour at room temperature. Coverslips were then washed with PBS for 3 x 5-minute washes, followed by incubation with secondary antibodies diluted in blocking solution for 1 hour at room temperature. For experiments requiring nuclear labeling, Hoechst dye 3342 (Molecular Probes, H3570) was included during the secondary antibody incubation at a final concentration of 0.5 µg/mL. Coverslips were then washed 3 x 5 minutes in PBS, rinsed briefly in ddH₂O, and mounted in ProLong Gold Antifade Mountant (Thermo Fisher Scientific, P36930). All steps of this procedure were performed with the samples protected from ambient light.

### Selection of Astrocytes

We have previously shown that astrocytes cocultured with neurons progressively undergo stellation within one week of coculture (1). Within this system, astrocytes can display a range of morphological complexity that correlates with protein expression; for example GLAST and AQP4 are preferentially enriched in highly branched astrocytes (Fig. S1C-D). We therefore have applied the following criteria to select astrocytes for analysis. For early-stage branch analyses (DIV3-4), we selected astrocytes with radial symmetry, 3-4 thick primary branches, and rudimentary secondary branchlet formation. For later-stage analyses (DIV6-10), astrocytes were selected based on radial symmetry, the presence of at least 3-4 primary branches (typically more), non-overlapping territories with neighboring astrocytes, and a high degree of arborization, including well-defined tertiary and quaternary branchlets.

### Validation of Coculture

To validate the purity of the neuron preparation (Fig. S1B), non-transgenic cortical neurons were maintained for 8 DIV and immunostained as described above. Z-stacks were obtained spanning the entire depth of the coculture at 0.2 µm intervals. Maximum intensity projections of z-stacks were generated in Fiji. Cell nuclei labeled with Hoechst were used to count viable cells using the ‘Cell Counter’ tool. Nuclei counts were overlaid onto images with cell-type-specific markers to determine the proportion of cells positive for each marker.

To validate the purity of the glia in coculture with neurons (Fig. S1D-E), non-transgenic cortical neurons and GFP-LC3 transgenic astrocytes were cocultured for 7 DIV and immunostained as described above. Z-stacks were obtained spanning the entire depth of the coculture at 0.2 µm intervals. To assess the purity of the astrocyte population of interest in this study, fields of view were selected based on the presence of astrocytes with stellate morphology. Maximum intensity projections of z-stacks were generated in Fiji. GFP(LC3) signal was used to identify and count glia within each field using the Cell Counter tool. Counts were overlaid onto images with cell-type-specific markers to determine the proportion of cells positive for each marker.

### Pharmacological Manipulation of TRPML1 Activity

#### Preparation and validation of TRPML1 modulators

ML-SA1 and ML-SI3 were dissolved in DMSO at stock concentrations of 20 mM and 10 mM, respectively. As both compounds have limited stability in solution, stocks were stored at −80°C and used within 1-2 months. Whenever possible, the efficacy of ML-SA1 and ML-SI3 was validated by on-scope addition to measure changes in GCaMP6-TRPML1 fluorescence intensity, as demonstrated in Fig. 2D.

**Figure 2.**
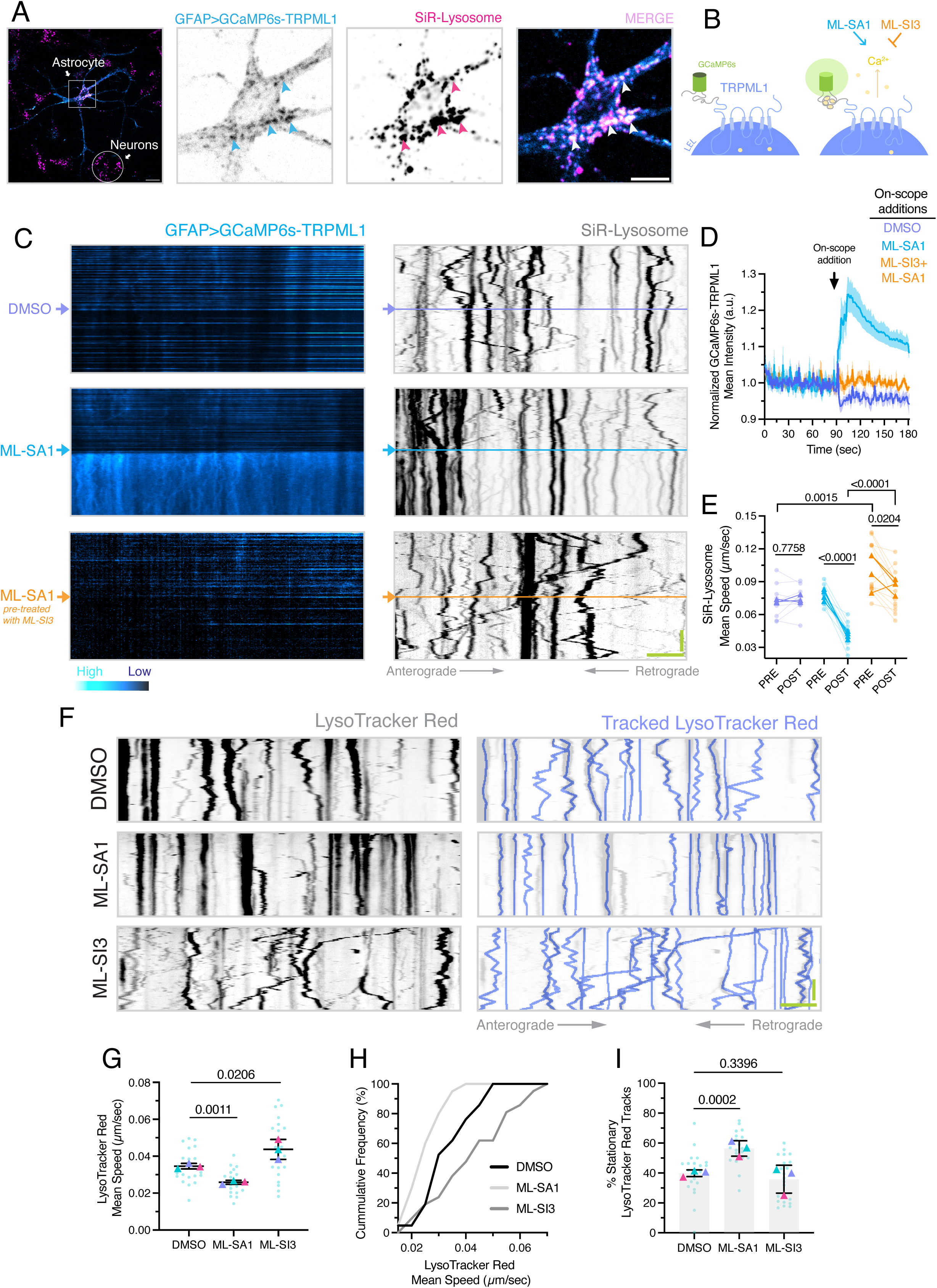
TRPML1 activation dampens LEL motility in astrocytic branches. **(A)** Representative single-plane image of a non-transgenic DIV7 cocultured astrocyte transduced with lentiviral GCaMP6s-TRPML1 (expressed under a shortened GFAP promoter) and co-labeled SiR-Lysosome; scale bar, 20 µm. Box denotes region of zoom-in of GCaMP6s-TRPML1 and SiR-Lysosome; scale bar, 10 µm. LELs co-positive for GCaMP6s-TRPML1 and SiR-Lysosome are denoted with closed arrowheads. **(B)** Schematic of GCaMP6s-TRPML1 fusion protein; TRPML1 forms a homotetrameric channel on LEL membranes (82). Noted are pharmacological tools used to modulate TRPML1 activity. **(C-E) (C)** Representative kymographs of GCaMP6s-TRPML1 and SiR-Lysosome along the astrocyte branch; GCaMP6s-TRPML1 was used as a space-fill to delineate the branch boundary. Arrows indicate the time of on-scope addition of DMSO (solvent control), ML-SA1 (60 µM), or co-addition of ML-SA1 (60 µM) and ML-SI3 (30 µM). Prior to imaging, astrocytes were pretreated for 1.5 hours with DMSO (paired with the on-scope addition of DMSO or ML-SA1) or ML-SI3 (30 µM; paired with the on-scope co-addition of ML-SA1 and ML-SI3); SiR-Lysosome was added during the final 30 minutes of pretreatment. Given that GCaMP6s-TRPML1 overexpression alone dampens LEL motility, imaging was performed at coculture DIV5, when baseline motility is higher and affords a greater dynamic range for detecting motility changes. Horizontal bar, 5 µm; vertical bar, 30 sec. **(D)** Quantification of the mean intensity of GCaMP6s-TRPML1 along the astrocyte branch, normalized to the average mean intensity before on-scope addition. Shown are means ± SEM; N=9-12 branches (one branch per astrocyte) across 3-4 biological replicates. **(E)** Quantification of SiR-Lysosome mean speed based on kymograph analysis. Small, transparent paired points indicate the average SiR-Lysosome mean speed per branch pre-versus post-treatment; large, opaque paired points indicate the biological replicate mean pre-versus post-treatment. Shown are p-values from a mixed-effects model with Tukey’s post-hoc correction applied to biological replicate values; N=9-12 branches (one branch per astrocyte) across 3-4 biological replicates. **(F-I) (F)** Kymographs of LysoTracker Red motility along a primary-to-secondary astrocytic branch; the GFP-LC3 transgene was used as a space-fill to delineate the branch boundary. Cocultures were treated for 30 minutes with DMSO (solvent control), ML-SA1 (60 µM), or ML-SI3 (30 µM) prior to live-cell imaging. LysoTracker Red puncta that were manually tracked (shown in periwinkle) are overlaid onto the kymographs. Horizontal bar, 5 µm; vertical bar, 30 sec. All quantification is derived from the kymograph analysis**: (G)** mean speed of LysoTracker Red tracks, **(H)** cumulative frequency of LysoTracker Red mean speed (derived from the technical replicates in G), and **(I)** percentage of stationary (or immobile) tracks. **(G, I)** Small dots indicate measurements from individual branches from individual astrocytes (e.g., technical replicates) from each of the independent experiments; larger triangles indicate the corresponding mean from each independent experiment (e.g., the biological replicates). Horizontal bars represent the mean of biological replicates ± SD. Shown are p-values from an LME model; N=20-21 branches (one branch per astrocyte) across 3 biological replicates; DIV7-8 of coculture.

#### Live Cell Imaging

For short-term drug treatments (Fig. 2F-I; Fig. 3F-I; Fig. 4C-J), cocultures were incubated for 30 minutes in coculture medium containing DMSO (0.3%), 60 µM ML-SA1, or 30 µM ML-SI3, along with SiR-Lysosome (1 µM) or LysoTracker Red (100 nM). Coverslips were then washed twice with HibE imaging medium and transferred to imaging chambers containing fresh HibE imaging medium containing the same drug treatment to prevent washout during imaging. Movies were acquired at 1 frame per second for 120 seconds; total imaging time per coverslip did not exceed 30-40 minutes.

**Figure 3.**
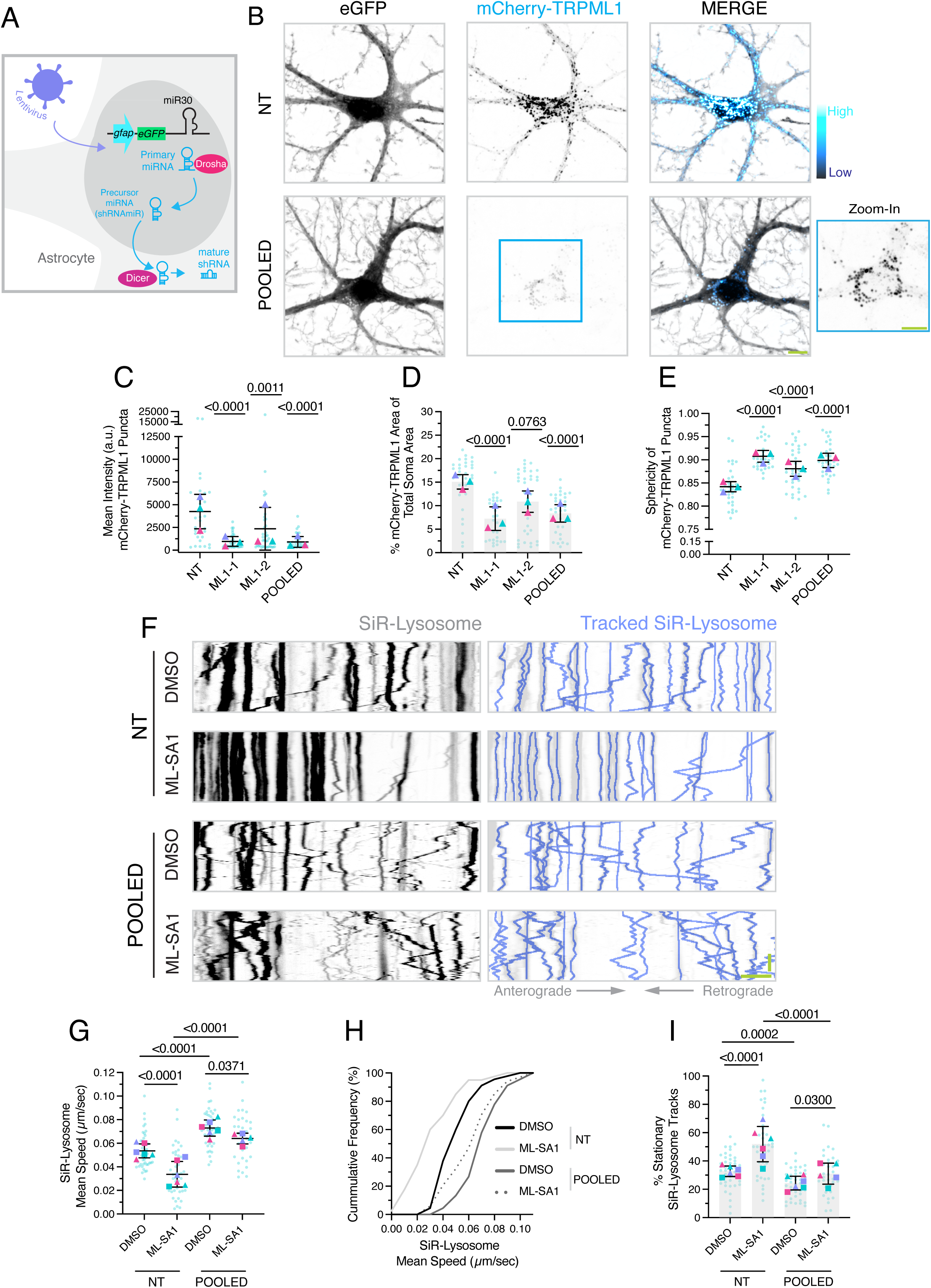
TRPML1 knockdown increases lysosome motility and antagonizes the ML-SA1-induced arrest of LEL motility along astrocyte branches. **(A)** Schematic of the miR30-shRNA system used to knockdown TRPML1 expression in cocultured astrocytes. shRNA sequences are embedded within a miR30 backbone to enable Drosha/Dicer-dependent processing from a Pol II-driven transcript, permitting astrocyte-specific knockdown when paired with a GFAP promoter. **(B-E) (B)** Representative maximum-intensity projections of z-stacks of DIV7 cocultured astrocytes co-expressing lentiviral eGFP (identifies astrocytes transduced with shRNAmiR) and mCherry-TRPML1 under a shortened GFAP promoter. The shRNA constructs included either a non-targeting shRNA control (denoted as NT) or a pool of two individual shRNAs targeting TRPML1 (denoted as pooled; individual shRNAs are denoted ML1-1 and ML1-2). mCherry-TRPML1 images are displayed with identical minimum and maximum intensity settings across treatment conditions; scale bar, 10 µm. Box indicates location of zoom-in of mCherry-TRPML1 in the knockdown condition. Zoom-in is adjusted individually to emphasize the sphericity of residual mCherry-TRPML1 puncta, consistent with lipid accumulation observed in MLIV (48, 49); scale bar, 10 µm. **(C)** Quantification of mCherry-TRPML1 puncta mean intensity. **(D)** Quantification of the percentage of total area occupied by mCherry-TRPML1 normalized to soma area. **(E)** Quantification of the sphericity of mCherry-TRPML1 puncta. **(C-E)** Horizontal bars represent the mean of biological replicates ± SD; shown are p-values from a LME model; N=29-35 astrocytes across 3 biological replicates; DIV7-10 of coculture. **(F-I) (F)** Kymographs of SiR-Lysosome motility along primary-to-secondary astrocytic branches; eGFP (shRNAmiR) was used as a space-fill to delineate the branch boundary. Cocultures transduced with either the non-targeting control or the pool of two individual TRPML1-targeting shRNAs were treated for 30 minutes with DMSO (solvent control) or ML-SA1 (60 µM) prior to imaging. SiR-Lysosome puncta that were manually tracked (shown in periwinkle) are overlaid onto kymographs. Horizontal bar, 5 µm; vertical bar, 30 sec. All quantification is derived from kymograph analysis: **(G)** mean speed of SiR-Lysosome tracks, **(H)** cumulative frequency of SiR-Lysosome mean speed (derived from the technical replicates in G), and **(I)** percentage of stationary (or immobile) tracks. **(G, I)** Horizontal bars represent the mean of biological replicates ± SD; shown are p-values from a LME model with Holm’s correction for multiple comparisons; N=34-45 branches (one branch per astrocyte) across 5-6 biological replicates; DIV6-8 of coculture.

**Figure 4.**
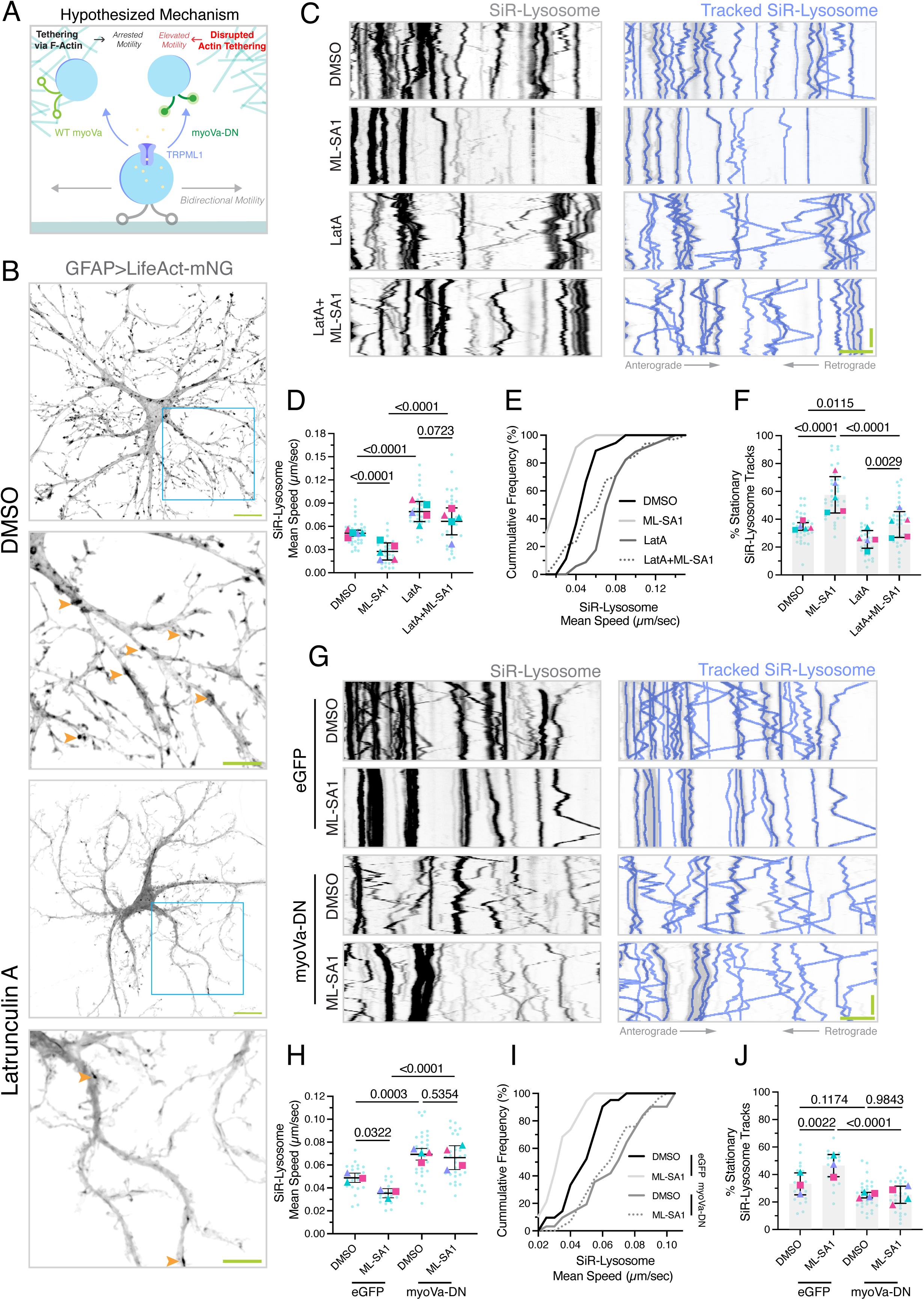
TRPML1-induced LEL motility arrest is dependent on F-actin and myosin-Va. **(A)** Schematic depicting the proposed model for TRPML1-induced LEL arrest. Motile LELs exhibit predominately bidirectional trajectories. Tethering to F-actin via myosin-Va may reduce LEL motility. Perturbations to F-actin or myosin-Va activity are hypothesized to disrupt LEL tethering and increase LEL motility. **(B)** Representative maximum-intensity projections of z-stacks acquired during live-cell imaging of non-transgenic cocultured astrocytes transduced with lentiviral LifeAct-mNeonGreen (LifeAct-mNG; expressed under a shortened GFAP promoter) at DIV7. Cocultures were treated for 30 minutes with DMSO (solvent control) or Latrunculin A (LatA; 5 µM). Cyan box denotes the area shown in the zoom-in. Orange arrowheads denote hot spots or patches of actin along the shaft of the branch, in filopodia, and in small lamellar structures. Scale bars, 20 µm (main) and 10 µm (zoom-in). **(C-F) (C)** Kymographs of SiR-Lysosome motility along astrocytic branches; the GFP-LC3 transgene was used as a space-fill to delineate the branch boundary. Cocultures were treated for 30 minutes with DMSO (solvent control), ML-SA1 (60 µM), LatA (5 µM), or a co-treatment of ML-SA1 and LatA. SiR-Lysosome puncta that were manually tracked (shown in periwinkle) are overlaid onto kymographs. Horizontal bar, 5 µm; vertical bar, 30 sec. All quantification is derived from kymograph analysis: **(D)** mean speed of SiR-Lysosome tracks, **(E)** cumulative frequency of SiR-Lysosome mean speed (derived from the technical replicates in D), and **(F)** percentage of stationary (or immobile) tracks**. (D, F)** Horizontal bars represent the mean of biological replicates ± SD; shown are p-values from a LME model with Holm’s correction for multiple comparisons; N=34-36 branches (one branch per astrocyte) across 5 biological replicates; DIV7-8 of coculture. **(G-J) (G)** Kymographs of SiR-Lysosome motility along astrocytic branches expressing lentiviral eGFP or myosin-Va-DN (C-terminal tail domain fused to the C-terminus of eGFP) under a shortened GFAP promoter. Cocultures were treated for 30 minutes with DMSO (solvent control) or ML-SA1 (60 µM). SiR-Lysosome puncta that were manually tracked (shown in periwinkle) are overlaid onto kymographs. Horizontal bar, 5 µm; vertical bar, 30 sec. All quantification is derived from kymograph analysis: **(H)** mean speed of SiR-Lysosome tracks, **(I)** cumulative frequency of SiR-Lysosome mean speed (derived from the technical replicates in H), and **(J)** percentage of stationary (or immobile) tracks**. (H, J)** Horizontal bars represent the mean of biological replicates ± SD; shown are p-values from a LME model with Holm’s correction for multiple comparisons; N=19-29 branches (one branch per astrocyte) across 3-4 biological replicates; DIV6-7 of coculture.

For extended drug treatments (Fig. 1E-K and L-N, Fig. S2F-G; Fig. S4A-H), cocultures were incubated for 2 hours in coculture medium containing DMSO (0.1%), 20 µM ML-SA1, or 10 µM ML-SI3; LysoTracker Red (100 nM) and/or SiR-Lysosome (1 µM) were included in the final 30 minutes. Coverslips were then washed twice with HibE imaging medium and transferred to imaging chambers containing fresh HibE imaging medium containing the same drug treatment to prevent washout during imaging. Movies were acquired at 1 frame per second for 120 seconds; only two replicates in Fig. 1E-G and Fig. S2F-G, and one replicate in Fig. 1J-K and Fig. S4A-D, were acquired for 60 seconds rather than 120 seconds. Total imaging time per coverslip did not exceed 30-40 minutes.

For on-scope delivery experiments (Fig. 2C-E), cocultured astrocytes at DIV5 were transduced with GCaMP6s-TRPML1 under a shortened GFAP promoter. We noticed that GCaMP6s-TRPML1 overexpression alone can dampen LEL motility. Thus, imaging was performed at coculture DIV5, when baseline motility is higher and affords a greater dynamic range for detecting motility changes. Coverslips were incubated for 1.5 hours in coculture medium supplemented with DMSO (0.3%) or ML-SI3 (30 µM); SiR-Lysosome (1 µM) was included in the final 30 minutes. Coverslips were then washed twice with HibE imaging medium and transferred to imaging chambers containing fresh HibE imaging medium containing the same drug treatment to prevent washout during imaging. Samples pre-treated with DMSO received an on-scope delivery of HibE imaging medium containing DMSO (final concentration of 0.3%) or ML-SA1 (final concentration of 60 µM). Samples pre-treated with ML-SI3 received an on-scope delivery of HibE imaging medium containing ML-SI3 and ML-SA1 to achieve a final concentration of 30 µM and 60 µM, respectively. Movies were acquired at 1 frame per second for 180 seconds; on-scope deliveries were performed between frames 89 and 90.

#### Immunofluorescence

Non-transgenic cocultures were treated with DMSO (0.3%), ML-SA1 (60 µM), or ML-SI3 (30 µM) in pre-conditioned coculture media for either 5 minutes (Fig. 5C-Fi) or for a time course of 5, 30, 60, and 120 minutes (Fig. S7I-K). Samples were then fixed and processed for immunostain as described above.

**Figure 5.**
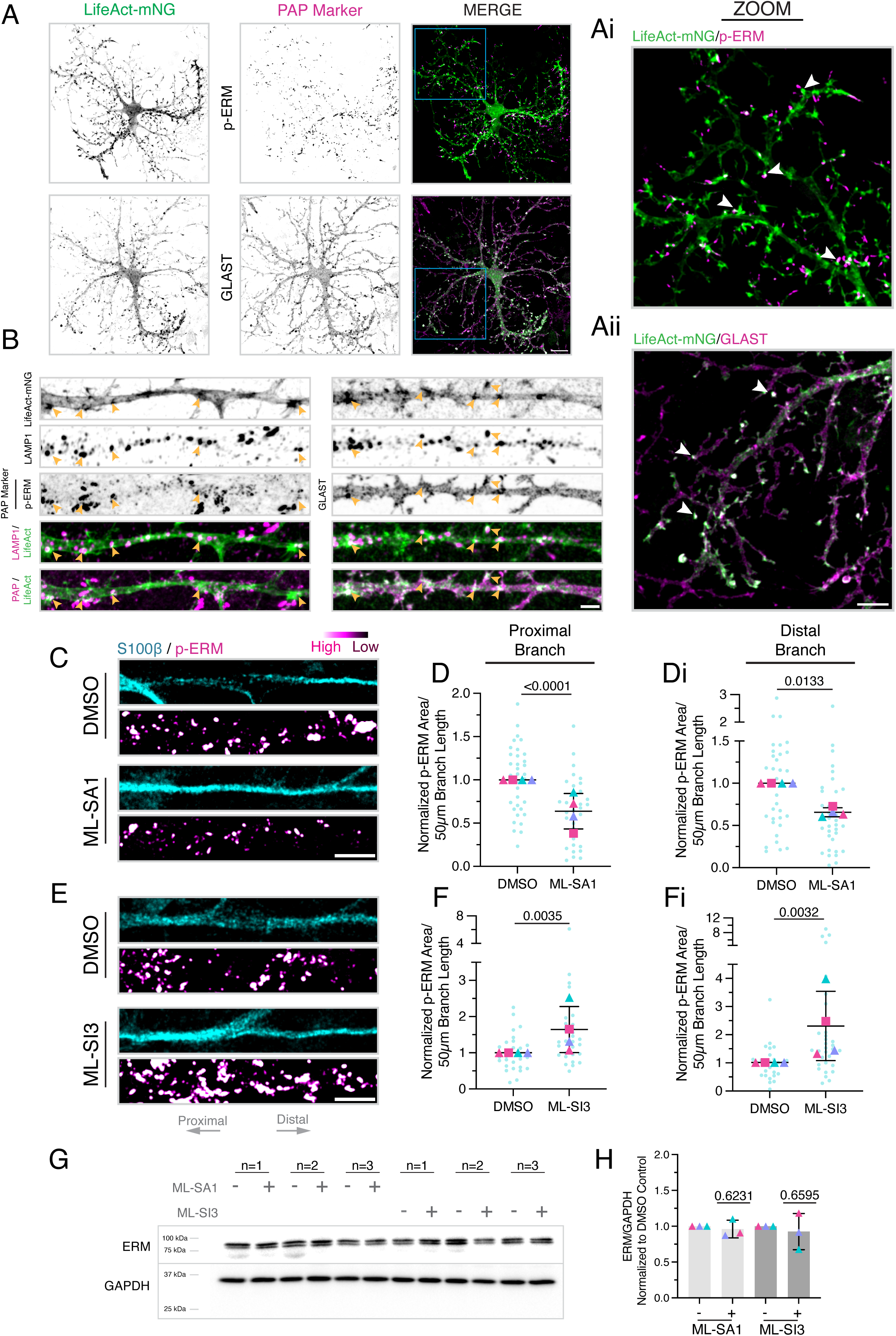
TRPML1 activity modulates the phosphorylation of PAP-enriched actin-membrane linkers ezrin-radixin-moesin. **(A)** Maximum-intensity projections of z-stacks acquired of DIV7 non-transgenic cocultured astrocytes transduced with lentiviral LifeAct-mNeonGreen (mNG) to visualize F-actin. Cocultures were immunostained for p-ERM proteins (phospho-ezrin Thr567, phospho-radixin Thr564, and phospho-moesin Thr558) or the plasma membrane marker GLAST/EAAT1. Scale bar, 20 µm. Boxed region denotes location of zoom-ins. **(Ai-ii)** Corresponding zoom-ins of astrocytic branches. White arrows denote F-actin-enriched foci co-positive for PAP-enriched markers. Scale bar, 10 µm. **(B)** Straightened branches from maximum-intensity projections of z-stacks acquired of DIV8 non-transgenic cocultured astrocytes transduced with lentiviral LifeAct-mNeonGreen (mNG) to visualize F-actin. Cocultures were immunostained for PAP-enriched markers p-ERM or GLAST and LEL-associated LAMP1. Orange arrowheads denote LAMP1-positive organelles positioned proximal to F-actin-enriched foci and PAP markers. Scale bar, 5 µm. **(C-Di) (C)** Straightened branches from maximum-intensity projections of z-stacks acquired of DIV7-8 non-transgenic cocultured astrocytes identified by immunostain for S100β and co-labeled for p-ERM. Cocultures were treated with DMSO (solvent control) or ML-SA1 (60 µM) for 5 minutes. S100β and p-ERM channels are each displayed with identical minimum and maximum intensity settings across treatment conditions. Scale bar, 10 µm. **(D-Di)** Corresponding quantification of the total area occupied by p-ERM-positive structures within the **(D)** proximal or **(Di)** distal 50 µm segment of the branch; values were normalized to the mean of the DMSO control per biological replicate. Horizontal bars represent the mean of biological replicates ± SD; shown are p-values from a LME model; N=34 branches (one branch per astrocyte) across 4 biological replicates. **(E-Fi) (E)** Straightened branches from maximum-intensity projections of z-stacks acquired of DIV7-8 non-transgenic cocultured astrocytes identified by immunostain for S100β and co-labeled for p-ERM. Cocultures were treated with DMSO (solvent control) or ML-SI3 (30 µM) for 5 minutes. S100β and p-ERM channels are each displayed with identical minimum and maximum intensity settings across treatment conditions. Scale bar, 10 µm. **(F-Fi)** Corresponding quantification of the total area occupied by p-ERM-positive structures within the **(F)** proximal or **(Fi)** distal 50 µm segment of the branch; values were normalized to the mean of the DMSO control per biological replicate. Horizontal bars represent the mean of biological replicates ± SD; shown are p-values from a LME model; N=28-31 branches (one branch per astrocyte) across 4 biological replicates. **(G-H) (G)** Immunoblot analysis of lysates from non-transgenic monocultured astrocytes (DIV6-7) treated with DMSO (solvent control), ML-SA1 (60 µM), or ML-SI3 (30 µM) for 5 minutes. Samples are immunoblotted for total ERM proteins (appear as doublets) and GAPDH (loading control). Shown are three independent biological replicates (denoted as n=1-3). **(H)** Corresponding densitometric analysis of total ERM levels normalized to GAPDH, and reported as a fold change relative to the DMSO control. Horizontal bars represent the mean of biological replicates ± SD; shown are p-values from a one-sample t-test; N=3 biological replicates.

#### Immunoblotting

For Fig. 5G-H and Fig. S7A-H, monocultured non-transgenic cortical astrocytes were treated with DMSO (0.3%), ML-SA1 (60 µM), MK6-83 (30 µM), or ML-SI3 (30 µM) in pre-conditioned glial media for 5 minutes prior to lysis, described below.

### Astrocyte-Specific Knockdown of TRPML1

For measurements of lysosomal motility (Fig. 3F-I), non-transgenic cocultures were transduced with lentiviral shRNAmiRs (coexpressed with eGFP) at a 1:10 dilution on coculture DIV2-4 and analyzed by live-cell imaging on coculture DIV6-8, for a total of 3-5 days of expression. For validation of knockdown efficiency (Fig. 3B-E), non-transgenic cocultures were co-transduced with shRNAmiRs and mCherry-TRPML1, each at a 1:10 dilution on coculture DIV2-6 and analyzed by live-cell imaging on coculture DIV6-10, for a total of 4 days of expression.

### Pharmacological Manipulation of F-Actin

#### Validation of Latrunculin-A

Non-transgenic cocultures at DIV4 of coculture were transduced with lentiviral LifeAct-mNeonGreen under a shortened GFAP promoter and imaged at DIV7 (Fig. 4B). Prior to imaging, cocultures were treated for 30 minutes in coculture medium supplemented with DMSO (0.25%) or Latrunculin-A (5 µM). Coverslips were washed twice with HibE imaging medium and transferred to imaging chambers containing fresh HibE imaging medium with the matching drug treatment to prevent washout during live-cell imaging. Z-stacks were acquired spanning the entire depth of the coculture at 0.2 µm z-intervals. Total imaging time per coverslip did not exceed 30-40 minutes.

#### Latrunculin-A and LEL Motility

Primary cortical neuron-astrocyte cocultures were prepared at DIV7-8; astrocytes were distinguished by the GFP-LC3 transgene and cocultured with non-transgenic neurons (Fig. 4C-F). Prior to imaging, cocultures were treated for 30 minutes in coculture medium supplemented with DMSO (0.3%), ML-SA1 (60 µM), Latrunculin-A (5 µM), or co-treatment with ML-SA1 (60 µM) and LatA (5 µM); SiR-Lysosome was also included at 1 µM final concentration. Coverslips were washed twice with HibE imaging medium and transferred to imaging chambers containing fresh HibE imaging medium with the matching drug treatment to prevent washout during live-cell imaging. Movies were acquired at 1 frame per second for 120 seconds; total imaging time per coverslip did not exceed 30-40 minutes.

### Myosin-Va Dominant Negative

Non-transgenic cocultures at DIV2-4 were transduced with lentiviral eGFP or a dominant negative myosin-Va construct (labeled myosin-Va DN) consisting of the C-terminal tail domain of myosin-Va fused to the C-terminus of eGFP. Each construct was expressed under a shortened GFAP promoter at a 1:10 dilution and analyzed by live-cell imaging at coculture DIV6-7, for a total of 3-4 days of expression (Fig. 4G-J).

### Pharmacological Manipulation of Synaptic Activity and Fluo-4 AM Imaging

In Fig. S6A-D, DIV7 cocultures of GFP-LC3 transgenic astrocytes and non-transgenic neurons were treated for 30 minutes in coculture medium supplemented with 50µM CNQX and 50µM AP5, or an equivalent volume of DMSO (0.2%) as a solvent control, at 37°C in a 5% CO_2_ incubator. Treatments also included SiR-Lysosome (1µM) and LysoTracker Red (100nM) to label LELs. Coverslips were then washed twice in HibE imaging medium and transferred to imaging chambers containing fresh HibE imaging medium with matching drug treatments to prevent washout. Movies were acquired at 1 frame per second for 120 seconds; total imaging time per coverslip did not exceed 30-40 minutes.

In Fig. S5D-Di, DIV7 cocultures of non-transgenic astrocytes and neurons were incubated with 4 µM Fluo-4 AM for 20 minutes in coculture medium at 37°C in a 5% CO_2_ incubator. Coverslips were then washed twice in HibE imaging medium and transferred to imaging chambers containing fresh HibE imaging medium. Movies were acquired at 1 frame per second for a total of 105 seconds. Between frames 29-30, 4AP and Bicuculline (diluted in HibE imaging medium) were added at a final concentration of 50µM 4AP and 50µM Bicuculline (S5D-Di). Between frames 89-90, CNQX and AP5 (diluted in HibE imaging medium) were added at a final concentration of 50µM CNQX and 50µM AP5 (Fig. S5D).

### Calcium Traces

#### GCaMP6s-TRPML1

Branch ROIs used for kymograph analysis of SiR-Lysosome described above (Fig. 2C-E) were also used to measure mean GCaMP6s-TRPML1 fluorescence intensity along astrocytic branches (line width=15 pixels). Mean fluorescence intensity was measured across all frames using the ‘Multi-Measure’ function in Fiji. Each frame was normalized to the mean intensity averaged over the baseline period prior to on-scope drug delivery (frames 1-89), and data were plotted as the mean across all measurements per condition in GraphPad Prism (70).

#### Fluo-4 AM

Non-transgenic cocultures at DIV7 were treated with Fluo-4 AM as described above. A square ROI was drawn over the entire neuronal meshwork to measure mean Fluo-4 AM fluorescence intensity. Mean fluorescence intensity was measured across all frames using the ‘Multi-Measure’ function in Fiji. Each frame was normalized to the mean intensity averaged over the baseline period prior to on-scope drug delivery (frames 1-29) and plotted in GraphPad Prism (70).

### BSA and DQ-BSA labeling

At DIV3 or DIV7-8, cocultures of non-transgenic neurons and GFP-LC3 transgenic astrocytes were co-incubated with 25 µg/mL BSA-Alexa 647 and 25 µg/mL DQ-Red-BSA diluted in pre-conditioned coculture medium for 2 hours, rinsed twice with HibE imaging medium, and imaged in fresh HibE imaging medium (Fig. 1A, C-D; Fig. S2A-B). Z-stacks were acquired spanning the entire depth of the coculture at 0.2 µm intervals.

### Kymograph Analysis for LEL Motility

Kymographs were generated in Fiji using the KymoToolBox plugin (PMID: 30044968; https://github.com/fabricecordelieres/IJ-Plugin_KymoToolBox) (71). Segmented lines (15 pixels in width) were drawn along the trajectory of a primary-to-secondary branch, beginning ∼10 µm from the soma and extending distally, using a fluorescence cell-fill marker as a guide. For Fig. 1E-G, 1I-K, 2F-I, 4C-F, S2F-G, S4A-D, S6A-D, the GFP-LC3 transgene served as a space fill marker in astrocytes. For all other datasets, eGFP (Fig. 1L-N, 3F-I, 4G-J, S4E-H) or GCaMP6s-TRPML1 (Fig. 2C-E) were used to delineate astrocyte branch morphology. For on-scope addition experiments in Fig. 2C-E, time-lapse movies were divided into pre-addition (frames 1-89) and post-addition (frames 90-180) segments before making kymographs. If a selected branch passed through regions densely populated by neuronal cell bodies or processes, the line was redrawn to exclude these areas and isolate the astrocytic branch signal. Kymographs ranged from ∼75-150 µm in length.

Our analysis was performed blinded to the sample identity. We used a Fiji macro that randomizes the kymographs in the dataset and assigns a numerical title to each image (the identity of the data was decoded only after the analysis was completed). LEL trajectories were manually traced with high fidelity using the segmented line tool (line width of 3 pixels). For LELs labeled with SiR-Lysosome, or LysoTracker Red, we only tracked vesicles whose entire trajectory was discernible throughout the entire kymograph; this criterion minimized signal from neighboring neurites (Fig. 1N). Indeed, nearly all SiR-Lysosome tracks were co-positive for LAMP1-mCherry, which was expressed exclusively in astrocytes, validating our method for selective identification of LELs in astrocytes (Fig. 1N).

Using the KymoToolBox plugin, we calculated mean vesicle speed (µm/sec) for each individual track. Since the high density of LELs in branches could confound individual track assignments, we averaged the mean speeds across all tracks measured per astrocyte branch; approximately 25-60 vesicles were tracked per branch. Individual vesicle tracks were classified as stationary if their cumulative distance travelled was less than 2 µm within the two-minute imaging window. Individual tracks were classified as retrograde- or anterograde-biased if movement in that direction comprised more than 55% of the total track duration. Non-stationary tracks that did not exhibit this directional bias and traveled >2 µm cumulative distance within the two-minute imaging window, were classified as bidirectional.

### Percentage Overlap between SiR-Lysosome and other LEL Markers

Kymographs of SiR-Lysosome were generated, blinded, and analyzed as described above. Kymographs were unblinded and SiR-Lysosome-positive tracks were overlaid onto the corresponding LAMP1-mCherry (Fig. 1M-N; Fig. S4E-H) or LysoTracker Red (Fig. 1E-G; Fig. S2F-G) kymograph from the same astrocytic branch. SiR-Lysosome-positive tracks that co-migrated with LAMP1-mCherry or LysoTracker Red tracks were classified as co-positive, and those lacking overlap were classified as colocalization-negative. Because SiR-Lyso consistently exhibited higher fluorescence intensity and photostability, LysoTracker Red or LAMP1-mCherry tracks with faint or partial overlap were conservatively classified as co-positive. For Fig. 1G and S2F-G, the remaining LysoTracker Red tracks were subsequently tracked and classified as SiR-Lysosome-negative.

### Kymograph Analysis for EB3-mNeonGreen

Non-transgenic cocultures were transduced with EB3-mNeonGreen on coculture DIV1 (for imaging on DIV3-4) or DIV3-4 (for imaging on DIV7-8) for a total of 2-3 days of expression prior to live-cell imaging. Movies were acquired at 1 frame per second for 120 seconds. Kymographs were generated by drawing a segmented line (15 pixels in width) along the trajectory of a primary-to-secondary branch, beginning ∼10 µm from the soma and extending distally, using EB3-mNeonGreen to delineate branch morphology. Analysis was performed blinded to the sample identity, as described above. EB3 comet trajectories were manually traced in Fiji using the segmented line tool (line width = 1 pixel) and quantified using the KymoToolBox plugin to get directional bias and comet density normalized to kymograph length. To capture branch-to-branch variability, we analyzed two branches per astrocyte. Reported values for technical replicates in Fig. S3D, F, H represent the mean of the two branches per cell. Reported values for technical replicates in Fig. S3E, G, I are the individual measurements from each branch.

### Ilastik Analysis of LEL Markers

#### DQ-BSA Red and BSA-647

Cocultures of GFP-LC3 transgenic astrocytes and non-transgenic neurons were incubated with BSA conjugates as described above prior to live-cell imaging. Z-stacks were acquired spanning the entire depth of the coculture at 0.2 µm intervals. For branch analyses (Fig. 1A, C-D; Fig. S2A-B), maximum intensity projections of z-stacks were generated in Fiji. Astrocytic branches were traced using the segmented line tool (line width = 60 pixels) along the trajectory of a primary to secondary branch, beginning ∼10 µm from the soma and extending distally, using GFP-LC3 to delineate branch morphology. Regions overlapping with dense patches of neuronal cell bodies were excluded from the analysis. Branch ROIs were straightened using the ‘Straighten’ function across channels (e.g., GFP-LC3, DQ-BSA-Red, and BSA-647) and then cross-sectional branch area was measured in the GFP-LC3 channel using the wand tracing tool. The branch boundary was overlaid onto channels of interest and signal outside the branch was cleared. Ilastik was used to identify and segment DQ-BSA-Red and BSA-647 puncta within the branch boundary. Total puncta area was quantified using the ‘Analyze Particles’ function in Fiji. The total area occupied by BSA-positive puncta for each marker was normalized to branch area and expressed as a percentage. To quantify colocalization between DQ-BSA-Red and BSA-647, the overlapping area between their respective ilastik segmentation masks was calculated using the ‘AND’ function in the Image Calculator tool. This overlapping area was then normalized to total BSA-647 area and expressed as a percentage.

#### mCherry-TRPML1

Non-transgenic cocultured astrocytes were transduced with eGFP (co-expressed with shRNAmiRs) and mCherry-TRPML1 both under a truncated GFAP promoter as described above (Fig. 3B-E). Z-stacks were acquired that spanned the entire depth of the coculture at 0.2 µm intervals. Maximum intensity projections of z-stacks were generated in Fiji. Soma boundaries were manually outlined using the ‘freehand tool’ in Fiji, referencing only the eGFP signal (and blinded to other channels) as a space-filling marker to delineate each cell body; cross-sectional areas of the resulting ROIs were then measured. These outlines were overlaid onto the corresponding channels of interest (e.g., mCherry-TRPML1) and signal outside the soma boundary was cleared. Ilastik (72) was then used to identify and segment mCherry-TRPML1-positive puncta within each soma. Puncta area, mean intensity, and sphericity were quantified using the ‘Analyze Particles’ function in Fiji; total puncta area was normalized to respective soma area and expressed as a percentage.

#### SiR-Lysosome

GFP-LC3 transgenic astrocytes were maintained in a monoculture as described above for DIV5-6 prior to live-cell imaging (Fig. S2C-E). Z-stacks were acquired spanning the entire depth of the astrocytes at 0.2 µm intervals. Maximum intensity projections of z-stacks were generated in Fiji. Cell boundaries were manually outlined using the ‘freehand tool’, referencing only the GFP-LC3 signal (and blinded to other channels) as a space-filling marker to delineate the cell boundary; cross-sectional areas of the resulting ROIs were then measured. Ilastik was then used to identify and segment SiR-Lysosome-positive puncta as described above. Puncta area and maximum intensity were quantified using the ‘Analyze Particles’ function. Total puncta area was normalized to respective cell area and expressed as a percentage relative to the solvent control.

### Ilastik Analysis of p-ERM Area

Z-stacks spanning the entire depth of the cocultures were acquired at 0.2 µm intervals, and maximum intensity projections were generated in Fiji. Primary-to-secondary branch trajectories were delineated using S100β (Fig. 5C-Fi, Fig. S7I-K), GFAP (Fig. S7I-K), or the GFP-LC3 transgene (Fig. S5J-K) as reference markers and traced using the segmented line tool (line width = 60 pixels), without consulting signal in other channels. A wider line width was used to capture p-ERM-positive structures emanating from the branch shaft. Each branch was traced beginning ∼10 µm from the soma and extending distally, and branch ROIs were then straightened using the ‘Straighten’ function in Fiji.

Because p-ERM signal was often enriched in the proximal branch, each branch was divided into proximal and distal 50 µm segments for separate analysis. Ilastik was used to identify and segment p-ERM-positive structures within each branch segment. Separate Ilastik classifiers were trained for proximal and distal regions within each biological replicate. Ilastik segmentations were imported into Fiji and total p-ERM-positive puncta area within each 50 µm segment was measured using the ‘Analyze Particles’ function. For each biological replicate, total p-ERM-positive area was normalized to the mean value of the solvent control.

For Figure S5J-K, branches at DIV3 are substantially shorter than at DIV7 and were therefore analyzed across their total length without segmentation. To enable direct comparisons across time points, this approach was applied consistently to both DIV3 and DIV7 samples.

### Ilastik Analysis of Synapses

Z-stacks spanning the entire depth of non-transgenic cocultures were acquired at 0.2 µm intervals; the plane which captured the largest cross-sectional area of the astrocyte (labeled with GFAP) was selected for analysis. From this slice, primary-to-secondary branch trajectories were delineated using GFAP (Fig. S5A-C) and traced using the segmented line tool (line width = 60 pixels) while blinded to the signal in other channels. A wider line width was used to capture synaptic structures proximal to astrocyte branches. Each branch was traced beginning ∼10 µm from the soma and extending distally, and branch ROIs in all channels were then straightened using the ‘Straighten’ function in Fiji. Ilastik classifiers were trained individually for Homer and Bassoon to identify and segment punctate synaptic structures. For Homer, the classifier was trained to omit the bright cytosolic signal prominent at DIV3. Segmentations were imported into Fiji and total puncta area was measured using the ‘Analyze Particles’ function. For each biological replicate, total puncta area positive for Homer or Bassoon was normalized to branch length and then normalized to the mean value of DIV3 to plot fold effects.

### Patch Clamp Recording

Cocultured non-transgenic neurons were recorded at 10 days of coculture (a total neuron age of DIV14). Before recording, coverslips were gently transferred to a recording chamber and held by a nylon slice holder and continuously perfused with oxygenated artificial cerebrospinal fluid (ACSF) containing the following: 126 mM NaCl, 2.5 mM KCl, 1.2 mM MgSO_4_, 2.4 mM CaCl_2_, 25 mM NaHCO_3_, 1.4 mM NaH_2_PO_4_, 11 mM glucose, and 0.6 mM sodium L-ascorbate, and continuously bubbled with 95% O_2_ and 5% CO_2_. Neurons were visualized with a 40X water-immersion objective under an Olympus BX61WI upright microscope. Recording pipettes were made from borosilicate glass (GC210F-10; Harvard Apparatus) with a Flaming-Brown P-97 puller (Sutter Instruments; tip resistance 5-8 MΩ). The pipette solution contained the following: 120 mM K-gluconate, 10 mM NaCl, 1 mM CaCl2, 10 mM EGTA, 10 mM HEPES, 5 mM Mg-ATP, 0.5 mM Na-GTP, and 10 mM phosphocreatine. For each recorded neuron, membrane properties were evaluated using current injections under current-clamp mode. The sEPSCs and sIPSCs were recorded under voltage-clamp mode. Electrophysiological recordings were acquired by an EPC-10 amplifier with PULSE and analyzed with FITMASTER (HEKA Electronics).

### Immunoblotting

Cells were washed with PBS and lysed in RIPA buffer (50 mM Tris-HCl pH 7.4, 150 mM NaCl, 1% Triton X-100, 0.5% sodium deoxycholate [Thermo Fisher Scientific, BP349-100], 0.1% SDS [Thermo Fisher Scientific/Invitrogen, 15553-027], and 1X Halt Protease and Phosphatase Inhibitor Cocktail supplemented with 5 mM EDTA [Thermo 78442]) for 30 min on ice. Lysates were centrifuged at 17,000 × *g* for 15 min at 4°C and supernatants were collected for SDS-PAGE and transferred to Immobilon-P PVDF membranes (Millipore). Membranes were blocked in 5% milk in TBS-Igepal buffer (24.8 mM Tris-HCl pH 7.4, 137 mM NaCl, 2.7 mM KCl, 0.05% Igepal [Sigma, I3021]) for 30 min at room temperature, and incubated overnight at 4°C with primary antibodies diluted in 5% milk in TBS-Igepal buffer. Membranes were washed 3 x 20 minutes in HRP wash buffer (50 mM Tris-HCl pH 8.0, 150 mM NaCl, 0.1% BSA, 0.05% Igepal) and then incubated for 45 min at room temperature with HRP-conjugated secondary antibodies diluted in HRP wash buffer. Membranes were washed 3 x 20 minutes in HRP wash buffer and developed using SuperSignal West Pico Chemiluminescent Substrate (Thermo Fisher Scientific, PI34580) on a ChemiDoc MP Imaging System (BioRad, #12003154).

### Quantitation of Immunoblot

Densitometric analysis was performed using the Gel Analyzer tool in Fiji. Band intensity was measured by drawing rectangular ROIs that encompassed each individual band; for pERM and ERM, the doublet bands were quantified together in a single ROI. Plot profiles were generated from each ROI and band intensity was quantified as the area under the peak. Values for p-ERM and ERM were normalized to values for the GAPDH loading control from each corresponding lane to control for differences in sample loading. The p-ERM/ERM ratio was calculated from the GAPDH-normalized values. Ratios were then normalized to the DMSO control condition and expressed as a fold change.

### Figure Preparation and Statistical Analysis

All superplot graphs were generated in GraphPad Prism (70). Semi-transparent circles indicate the measurements from individual cells (e.g., the technical replicates) from each of the independent experiments. Large opaque triangles and squares indicate the mean of each of the independent experiments (e.g., the biological replicates); independent experiments are color-coded. Horizontal bars represent the mean of biological replicates ± SD. The definition and values of N are provided in the figure legends for each individual graph. All cumulative frequency graphs were generated in GraphPad Prism.

Histograms and density estimates shown in Fig. S3E, G, and I were generated in R (version 2025.05.0+496) using the ggplot2 package. To visualize shifts in microtubule polarity at DIV3 versus DIV7-8, individual branch measurements were pooled within each timepoint and plotted as probability density histograms with bin widths held constant across conditions. Counts were normalized to probability density such that the total area under each distribution equals 1, enabling direct comparison of distribution shapes. A Gaussian kernel density estimate was overlaid on each histogram to provide a smoothed representation of the underlying distribution.

Our data have a nested structure, with multiple cells (technical replicates) measured within each biological replicate (independent cell preparations from different mice) across treatment conditions. As cells within the same biological replicate will share common sources of variability (e.g., variability in cell preparations, culturing conditions, or animals), measurements taken within a biological replicate are not independent of one another. To account for this non-independence, we fit linear mixed-effects (LME) models with treatment as a fixed effect and biological replicate as a random effect, using the nlme package in R (version 2025.05.0+496). This approach allows us to assess treatment effects, while accounting for within-replicate correlation and between-replicate variability. Model residuals were inspected for homoscedasticity. For heteroscedastic data, model fit was improved by incorporating a varIdent variance structure (Fig. 2G, 3C, 3I, 4D, 5F-Fi, S4D, S5K).

GraphPad Prism was used to perform the following statistical tests: In Fig. 2E, where cell throughput was low (e.g., 3 cells per biological replicate), a Mixed-Effects Model with Tukey’s post hoc correction was performed with the biological replicate mean values in GraphPad Prism. For immunoblot analysis (Fig. 5H, S7B-D, S7F-H), band intensities were normalized to the solvent control within each independent experiment, such that the control value equaled 1.0. Treatment group means were then compared to this fixed reference value using a one-sample t-test in GraphPad Prism. All figures were assembled in Adobe Illustrator.

### Use of AI

Perplexity and Claude AI tools were used only during the editing process to improve clarity and readability of text in this manuscript. ChatGPT was used to aid in writing the code for statistical analysis in R. The authors carefully reviewed and edited any AI-generated edits and take full responsibility for the content of the publication.

## RESULTS

### Astrocytes cocultured with neurons can develop a highly branched architecture to investigate lysosome biology

The organization of the lysosomal network in astrocytes is poorly understood. Much of the work to date exploring lysosome biology in neural cell types has been performed in neurons. To determine the distribution and dynamics of lysosomal organelles in astrocytes, we used an astrocyte-neuron coculture model to better recapitulate *in vivo* astrocyte biology (1, 69, 73). In this system, primary mouse cortical neurons are cultured for 4-5 days *in vitro* (DIV) to develop a dense meshwork of neurites (Fig. S1A). We validated that the neuron preparation is highly enriched for neurons (∼96% pure) and largely depleted of glial cells (<5% of total cells) (Fig. S1B) (74). Primary mouse cortical glia enriched for astrocytes (74) are then plated on top of the neuronal meshwork and cocultured for DIV6-10 (neurons are a total of DIV11-14) (Fig. S1A). This staggered addition of astrocytes onto an established neurite meshwork is critical as direct contact and secreted factors from developed neurons promote astrocyte maturation (Fig. S1C) (75, 76).

We validated the purity and maturation state of the glial preparation using glial cells isolated from a transgenic mouse line expressing GFP-LC3 (LC3; microtubule-associated protein-1 light chain 3), in combination with immunostaining for a panel of glial-specific markers. LC3 is a ubiquitin-like protein that in its cytosolic form can delineate cellular morphology, and when lipidated, labels autophagosomes (68). Thus, the GFP-LC3 transgene enables us to identify glial cells in the coculture and their morphological development. Using GFP-LC3 transgenic glia, we validated that the glial preparation is highly enriched for astrocytes and is largely devoid of oligodendrocyte precursors, oligodendrocytes, and microglia (Fig. S1D-E). GFP-LC3 labeling further confirmed the transition from the polygonal morphology of monocultured astrocytes to the highly branched, stellate morphology exhibited by mature astrocytes in coculture (Fig. S1C-D). Indeed, Fig. S1C reveals the branched morphologies of astrocytes in coculture with a dense neuronal meshwork. Notably, astrocyte-specific plasma membrane markers (e.g., GLAST and AQP4) provide a more complete delineation of astrocyte morphology, including finer structural features (Fig. S1C-D). Combined, primary astrocytes cocultured with neurons acquire proteomic signatures and morphological features characteristic of astrocytes *in vivo*.

### Lysosomes concentrate in the soma and become less motile during astrocyte maturation in coculture with neurons

To define the lysosomal population in astrocyte branches, we performed live-cell imaging using a spinning disk confocal microscope of astrocytes at DIV3 versus DIV7-8 of coculture (Fig. 1A). These time points mark a significant transformation in astrocyte morphology: astrocytes analyzed at DIV3 formed rudimentary primary and secondary branch structures, whereas astrocytes selected at DIV7-8 exhibited a higher degree of branch arborization that includes tertiary and quaternary branchlets (Fig. 1A). To delineate astrocyte morphology during live-cell imaging, we cocultured GFP-LC3 transgenic astrocytes with non-transgenic neurons (Fig. 1A). We then employed a panel of markers to label various stages of organelle maturation along the endolysosomal spectrum (Fig. 1B). To distinguish proteolytically active compartments from total endocytic compartments, we incubated cocultures with fluorescently-labeled conjugates of BSA, a fluid phase marker that is internalized by endocytosis (Fig. 1B). BSA-Alexa Fluor 647 (abbreviated BSA-647) labels all endocytic compartments, while DQ-Red-BSA is a dye-quenched form of BSA that fluoresces only upon proteolytic cleavage, reporting mature lysosome activity (Fig. 1A-B).

At DIV3 of coculture, DQ-BSA-positive puncta were distributed throughout the soma, proximal and distal branches in astrocytes (Fig. 1A). However, by DIV7-8 of coculture, DQ-BSA-positive puncta were largely restricted to the soma and proximal branch (Fig. 1A). To quantify these changes, we used Ilastik, a machine learning program to identify and segment BSA puncta (72). We found a significant reduction in DQ-BSA puncta area within astrocytic branches from DIV3 to DIV7-8, but no corresponding change in BSA-647 (Fig. 1C, S2B). Since BSA-647 levels remained constant across time points, the reduction in DQ-BSA signal likely does not reflect decreased endocytosis. Rather, the fraction of BSA-647-positive endosomes co-positive for DQ-BSA decreased from DIV3 to DIV7-8 (Fig. 1D), indicating a specific decline in lysosomal proteolytic activity over this maturation window. Together, these data suggest that as astrocytes mature, proteolytic activity becomes concentrated in the soma and proximal branches, establishing a gradient of degradative capacity along the branch arbor (Fig. 1H).

To corroborate these findings, we stained cocultures with the cell-permeant probes LysoTracker Red and SiR-Lysosome. LysoTracker Red accumulates in acidic compartments (pH <∼6). SiR-Lysosome is a fluorescent conjugate of pepstatin A, a pentapeptide that binds with high affinity to the active site of Cathepsin D at acidic pH around 3.5-5 (77, 78). SiR-Lysosome is more selective for labeling late endosomes-lysosomes, an acidic environment that sustains the active conformation of Cathepsin D, as validated by Correlative Light and Electron Microscopy (79). To confirm probe specificity, we showed that SiR-Lysosome signal is reduced upon co-treatment with a protease inhibitor cocktail containing excess unlabeled pepstatin A, which competes with SiR-Lysosome for occupancy of the Cathepsin D active site (Fig. S2C-E). We then performed dual-color live-cell imaging of LysoTracker and SiR-Lysosome in GFP-LC3 transgenic astrocytes cocultured with non-transgenic neurons (Fig. 1E). We generated kymographs along GFP-LC3-positive branches and quantified the density of individual puncta and overlap between each marker (Fig. 1E). To exclude potential signal from neighboring neurites, we only tracked puncta that moved within the trajectory of the astrocyte branch throughout the entire kymograph.

We found that nearly all SiR-Lysosome puncta were co-positive for LysoTracker Red (Fig. S2F), consistent with the activation of lysosomal proteases within an acidic environment. Similar to our observations with DQ-BSA, the density of SiR-Lysosome puncta decreased from DIV3 to DIV7-8, while the density of LysoTracker puncta remained constant (Fig. 1E-F; S2G). At DIV3, ∼80% of LysoTracker puncta were co-positive for SiR-Lysosome and by DIV7-8, this fraction reduced to ∼62% (Fig. 1E, G). Thus, as astrocytic branches mature, the density of acidic organelles remains constant, but the percentage that are proteolytically competent decreases. Notably, SiR-Lysosome puncta remain detectable at DIV7-8, indicating some degree of proteolytic activity in more mature branches.

To further characterize lysosomal organelles in mature astrocyte branches, we co-transduced non-transgenic astrocyte-neuron cocultures with two lentiviral constructs driven by a truncated GFAP promoter to restrict expression to astrocytes: LAMP1-mCherry, a marker for late endosomes and lysosomes, and soluble eGFP, which provides a cytosolic volume fill of the branch (Fig. 1L). Cocultures were additionally labeled with SiR-Lysosome and imaged at DIV7-8 of coculture. LAMP1-positive organelles in mature astrocyte branches were largely co-positive for SiR-Lysosome, though a subset of LAMP1 puncta were SiR-Lysosome-negative (Fig. 1M), consistent with prior reports that LAMP1 labels a heterogeneous organelle population including non-degradative compartments (35–37). Using the reciprocal analysis, ∼82% of SiR-Lysosome puncta were co-positive for LAMP1-mCherry (Fig. 1M-N), further confirming that SiR-Lysosome labels LELs (Fig. 1N). Furthermore, the high degree of overlap between SiR-Lysosome and astrocyte-restricted LAMP1-mCherry confirms that our tracking methods detect SiR-Lysosome puncta specifically within astrocyte branches, and not overlapping neurites, with high fidelity (Fig. 1N).

In addition to their identity, we tracked changes in LEL dynamics during branch maturation by kymograph analysis. At DIV3, most SiR-Lysosome puncta were motile, exhibiting a combination of bidirectional (∼56%) and directional movement (∼22% anterograde, ∼16% retrograde), with few stationary puncta (∼7%) (Fig. 1I-J). By DIV7-8, SiR-Lysosome puncta remained predominantly bidirectional (∼42%), but the stationary fraction increased nearly 3.5-fold (∼25%) (Fig. 1I-J), and mean speed decreased significantly (Fig. 1K). Interestingly, this shift in LEL motility coincides with a reorganization of the microtubule cytoskeleton (Fig. S3). As astrocyte branches mature, the microtubule cytoskeleton transitions from a predominantly plus-end-out orientation at DIV3-4 to a more mixed polarity by DIV7-8, a shift that may promote simultaneous engagement of opposing motor forces and contribute to LEL stalling (Fig. S3). Together, these findings reveal that astrocyte maturation in coculture with neurons is accompanied by a spatial reorganization of LEL compartments. Highly degradative lysosomes concentrate in the soma, while LELs within branches become less motile and less proteolytically active.

### TRPML1 regulates the motility of late endosomes/lysosomes in astrocyte branches

We next investigated mechanisms that regulate LEL dynamics in astrocyte branches. We focused our attention on TRPML1, a calcium-permeable cation channel on late endosomes/lysosomes (80). TRPML1 is a key regulator of various aspects of lysosome biology, including lysosome transport and trafficking (54–57). However, most of these studies have been performed in neurons and non-neural cell types (54–57). The prominent astrocytic pathology in *MCOLN1-/-* mice (50–53) further motivated our investigation into how TRPML1 regulates LEL trafficking in astrocytes. To monitor TRPML1 activity in astrocytes, we fused GCaMP6s to the N-terminus of TRPML1 and transduced cocultured astrocytes with GCaMP6s-TRPML1 expressed under a truncated GFAP promoter (Fig. 2A). GCaMP6s is positioned on the cytoplasmic side of the lysosomal membrane, allowing detection of juxta-lysosomal calcium (Fig. 2B). As expected, GCaMP6s-TRPML1 localized predominantly to proteolytically active lysosomes, as evidenced by colocalization with SiR-Lysosome (Fig. 2A). To modulate TRPML1 activity, we used the well-established small molecule agonist ML-SA1, and antagonist ML-SI3 (81, 82) (Fig. 2B), delivered on-scope during live-cell imaging of astrocyte branches.

At baseline, GCaMP6s-TRPML1 exhibited calcium oscillations that traveled along the astrocyte branch in a wave-like fashion, a pattern consistent with the detection of ER-derived calcium in proximity to lysosomes (Fig. 2C). These calcium oscillations persisted after addition of the DMSO solvent control (0.3%), although their signal was reduced (Fig. 2C-D). On-scope addition of ML-SA1 evoked a rapid and robust increase in GCaMP6s-TRPML1 signal, often revealing vesicular tracks in branches (Fig. 2C-D). Pretreatment with ML-SI3 dampened baseline oscillations and blocked the ML-SA1-evoked calcium increase (Fig. 2C-D). These data validate that ML-SA1 and ML-SI3 effectively stimulate or inhibit TRPML1 activity, respectively, in astrocytes. Moreover, ML-SI3 pretreatment blocks agonist-evoked responses, confirming target specificity of both small-molecules.

Next, we assessed the effects of TRPML1 modulation on LEL motility in astrocyte branches by kymograph analysis. On-scope delivery of the DMSO solvent control had no effect on the motility of SiR-Lysosome-positive puncta in astrocyte branches, as measured by mean-speed (Fig. 2C, E). Strikingly, on-scope delivery of ML-SA1 (60 µM) induced a rapid arrest in LEL motility (Fig. 2C, E). Importantly, the effects of ML-SA1 on LEL motility were attenuated by pre-treatment with ML-SI3. Indeed, pretreatment with ML-SI3 (30 µM for 1.5 hours) increased the motility of SiR-Lysosome puncta at baseline, as measured by mean-speed, and mitigated the arrest in motility elicited by ML-SA1 (Fig. 2C, E). Combined, our data indicate that modulating TRPML1 activity can regulate LEL motility in astrocyte branches; stimulating TRPML1 dampens LEL motility and inhibiting TRPML1 increases LEL motility.

As overexpression of GCaMP6s-TRPML1 may potentiate the effects of ML-SA1 or ML-SI3, we next examined whether modulating endogenous TRPML1 activity similarly regulates LEL motility. Cocultures at DIV7-8 (not expressing exogenous GCaMP6s-TRPML1) were treated for 30 minutes with ML-SA1 (60 µM), ML-SI3 (30 µM), or DMSO, and LEL dynamics were tracked using LysoTracker Red as in Fig. 1E. ML-SA1 treatment dampened LEL motility, as evidenced by decreased mean speed and an increased fraction of stationary tracks (Fig. 2F-I). This effect was also observed at a lower concentration of ML-SA1 (20 µM, 2 hours) at both DIV3 and DIV7-8 (Fig. S4A-D). Moreover, ML-SA1 dampened the motility of puncta co-positive for SiR-Lysosome and genetically-encoded LAMP1-mCherry, confirming similar effects on multiple lysosomal markers present within astrocyte branches (Fig. S4E-H). By contrast, ML-SI3 treatment elicited the opposite effect as ML-SA1. ML-SI3 increased the motility of both LysoTracker-positive (Fig. 2F, G) and LAMP1-mCherry/SiR-Lysosome co-positive puncta (Fig. S4E-H), as evidenced by an increase in mean speed. Together, these data confirm that modulating endogenous TRPML1 activity regulates LEL motility in astrocyte branches, consistent with our findings from GCaMP6s-TRPML1-expressing astrocytes.

Because ML-SA1 can induce lysosomal calcium release above physiological levels, we sought an orthogonal approach to corroborate our results with ML-SI3 and inhibit physiological TRPML1 activity. We employed a genetic approach to knockdown TRPML1 expression in astrocytes, which also serves as a model for MLIV (44–46, 50). We generated lentiviruses to express shRNAs targeting TRPML1 (labeled ML1) or a non-targeting control (labeled NT). ML1 or NT shRNAs were embedded within a miR30 backbone under control of a truncated GFAP promoter, enabling astrocyte-restricted Drosha/Dicer-dependent processing from a Pol II transcript (83) (Fig. 3A). eGFP was co-expressed from the same construct to identify transduced astrocytes (Fig. 3A). As we were unable to find a commercially available antibody that reliably recognized mouse TRPML1, we validated the TRPML1 knockdown by co-transducing astrocytes with murine mCherry-TRPML1 and quantifying signal loss. In the NT control, mCherry-TRPML1 localized to puncta enriched in the soma and was found within astrocyte branches, consistent with the distribution of late endosomes and lysosomes (Fig. 3B). Two different ML1 shRNAs, applied individually or pooled, reduced mCherry-TRPML1 puncta intensity (ML1-1: ∼78% reduction, ML1-2: ∼45% reduction, pooled: ∼79% reduction) and area (ML1-1: ∼52% reduction; ML1-2: ∼28% reduction; pooled: ∼45% reduction) within the soma (Fig. 3C-D). Notably, residual mCherry-TRPML1 puncta in the ML1 samples exhibited increased sphericity, forming lipid droplet-like structures consistent with lysosomal lipid accumulation in MLIV (49), further validating knockdown efficacy (Fig. 3E).

We then analyzed changes in LEL dynamics by kymograph analysis (Fig. 3F). Knockdown of TRPML1 significantly enhanced the motility of LELs in astrocyte branches, as evidenced by an increase in the mean speed of SiR-Lysosome puncta and a decrease in the fraction of stationary tracks compared to the NT control (Fig. 3F-I). Thus, reducing endogenous levels of TRPML1 phenocopies our observations with the TRPML1 antagonist ML-SI3. As we report in Fig. 2F-I, ML-SA1 reduced the motility of LELs in astrocytes expressing the NT control (Fig. 3F-I). Importantly, TRPML1 knockdown antagonized the arrest in LEL motility caused by ML-SA1 (Fig. 3F-I). Thus, the effects of ML-SA1 on lysosome motility are via TRPML1 and not off-target. In total, we use multiple approaches to identify TRPML1 as a key regulator of LEL motility in astrocyte branches.

### TRPML1-dependent arrest in LEL motility involves myosin-Va tethering to the actin cytoskeleton

How does TRPML1 activation lead to an arrest in LEL motility in astrocyte branches? Clues may be drawn from studies in neurons. In response to synaptic activity, lysosomes can become positioned at the base of dendritic spines via mechanisms involving the actin cytoskeleton (2). Specifically, actin constrains lysosome motility in dendrites (2) and causes lysosomes to stall at F-actin patches located at the spine base (84). Notably, calcium release from mitochondria can promote their dissociation from microtubule tracks and enhance tethering to the actin cytoskeleton (85). Since TRPML1 activation triggers lysosomal calcium release (Fig. 2C), TRPML1 may similarly promote LEL association with actin in astrocytes (Fig. 4A). Thus, we examined the role of the actin cytoskeleton in TRPML1-mediated arrest of LELs in astrocyte branches.

We first visualized F-actin in mature cocultured astrocyte branches by expressing lentiviral LifeAct-mNeonGreen (mNG) under a truncated GFAP promoter. LifeAct is a 17 amino acid peptide derived from the actin-binding protein Abp140 of *Saccharomyces cerevisiae* that permits visualization of F-actin with minimal disruption to actin dynamics (86). Using live-cell imaging, we observed a range of actin structures along branches including hot spots or patches along the shaft of the branch, filopodia, and small lamellar structures (Fig. 4B). To assess the contributions of F-actin in LEL motility in coculture astrocytes, we disrupted F-actin using 5 µM Latrunculin-A (LatA) for 30 minutes. LatA is a cell-permeable toxin that sequesters actin monomers, severs F-actin filaments, and accelerates filament depolymerization (87, 88). Indeed, LatA treatment reduced the abundance of F-actin structures along the astrocyte branch (Fig. 4B).

By kymograph analysis, we found that LatA treatment (5 µM for 30 minutes) increased LEL motility, as evidenced by increased mean speed and a decreased fraction of stationary SiR-Lysosome puncta (Fig. 4C-F). Thus, F-actin is important for restricting LEL motility in astrocyte branches. We next asked whether the TRPML1-induced LEL motility arrest requires F-actin (Fig. 4A). As we showed previously, ML-SA1 alone led to an arrest in LEL motility in astrocytes (Fig. 4C-F). However, co-treatment of ML-SA1 with LatA strongly attenuated this dampening in motility, resulting in LEL mean speeds comparable to those of LatA alone (Fig. 4D-E). LatA and ML-SA1 co-treatment also resulted in a modest but significant increase in the fraction of stationary tracks relative to LatA alone, albeit this fraction was lower than ML-SA1 alone (Fig. 4F). Combined, these data suggest that the TRPML1-induced arrest in lysosome motility is dependent on an intact actin cytoskeleton.

The sustained effect of TRPML1-induced LEL stalling suggested a mechanism of anchoring to the actin cytoskeleton rather than passive entrapment. Interestingly, prior studies have identified myosin-Va in the tethering of LELs at actin-enriched patches within dendrites (84). Myosin-Va is an unconventional myosin capable of both processive organelle transport and static tethering to the actin cytoskeleton (89, 90). Indeed, myosin-Va functions as an actin-based tether for other lysosome-related organelles, including secretory granules (91), phagolysosomes (92), and melanosomes (93, 94). Myosin Va has also been implicated in the docking of synaptic vesicles (95), and the transport of ER into dendritic spines (96, 97). Together, these roles support myosin-Va as a versatile motor for positioning a variety of organelles at actin-rich sites. Notably, myosin-Va activity is regulated by calcium and calmodulin (98–100). Astrocytes express myosin-Va (101–103), but whether myosin-Va regulates LEL tethering in astrocyte branches has not been characterized.

To test whether myosin-Va mediates LEL arrest in response to TRPML1 activation (Fig. 4A), we transduced cocultured astrocytes with a dominant negative myosin-Va construct (labeled myoVa-DN) consisting of the C-terminal tail domain of myosin-Va fused to the C-terminus of eGFP, under control of a truncated GFAP promoter. By retaining cargo-binding capacity while lacking motor activity, myoVa-DN competes with endogenous myosin-Va for cargo binding, thereby disrupting actin-based tethering (63, 84, 104). We found that overexpression of myoVa-DN significantly increased LEL motility compared to a control expressing eGFP alone (Fig. 4G-J). Specifically, expression of myoVa-DN increased SiR-Lysosome puncta motility, as measured by an increase in mean speed relative to the eGFP control (Fig. 4G-J). Importantly, while ML-SA1 evoked LEL motility arrest in eGFP-expressing astrocytes, this effect was abolished by myoVa-DN overexpression (Fig. 4G-J). Thus, these data demonstrate that TRPML1 promotes LEL motility arrest through myosin-Va-dependent tethering to the actin cytoskeleton.

### TRPML1 activity modulates phosphorylation of actin-membrane linkers enriched in astrocyte PAPs

What is the functional significance of this arrest in lysosome motility? Precedent from neurons suggests LELs arrest at postsynaptic compartments to locally influence synaptic function and structural remodeling (1, 2, 6), raising the question of whether analogous mechanisms operate in astrocytes. In astrocytes, we find that TRPML1-dependent stalling of lysosomes in branches requires engagement with the actin cytoskeleton (Fig. 4). Notably, actin in astrocytes is enriched at PAPs, dynamic structures that can overlap with synapses (13, 20, 21, 105). Astrocytes lacking TRPML1 exhibit significant alterations in actin- and synapse-associated proteins (53). Moreover, TRPML1 repositions LELs and reorganizes actin to locally activate myosin IIA-dependent contractility in the rear of migrating immune cells (106–108), suggesting a conserved role for TRPML1 in coupling lysosome positioning with regulating the actin cytoskeleton. Consistent with this idea, recent findings from Festa et al. show that TRPML1-positive lysosomes in nascent oligodendrocyte processes may be positioned to promote morphological maturation by activating Rac1/PAK1-dependent actin polymerization (109). Together, these connections motivated us to investigate whether TRPML1-dependent lysosome arrest positions LELs to serve local functions at astrocyte PAPs.

To examine this possibility, we first assessed the synaptic maturity in the astrocyte-neuron coculture and confirmed that cocultured astrocytes formed PAP structures. Consistent with prior reports that astrocytes accelerate synaptogenesis (110), neurons formed a dense synaptic meshwork by DIV7 of coculture, as evidenced by an increase in density and juxtaposition of pre- and postsynaptic markers, Bassoon and Homer, respectively (Fig. S5A-C). Calcium imaging with Fluo-4 AM revealed spontaneous oscillatory activity in the neuronal network that was enhanced by 4-AP/bicuculline treatment (antagonists of voltage-gated potassium channels and inhibitory γ-aminobutyric acid A receptors, respectively) (Fig. S5D-Di) and dampened with CNQX/AP5 (antagonists of excitatory AMPA and NMDA receptors, respectively) (Fig. S5D). Whole-cell patch-clamp recordings of cocultured neurons confirmed action potential firing upon current injection (Fig. S5E-G) and the presence of both spontaneous excitatory and inhibitory postsynaptic currents (i.e., sEPSCs and sIPSCs, respectively) (Fig. S5H). Thus, neurons in coculture with astrocytes show functional synaptic connections.

To measure PAPs, we immunostained for p-ERM, the phosphorylated forms of ezrin, radixin, and moesin, which are structural components enriched in PAPs (21, 22, 105, 111). Within the mature brain, ezrin expression is largely restricted to astrocytes (21, 22, 111). ERM proteins link the plasma membrane to F-actin to control membrane dynamics and tension, and their activity is regulated by phosphorylation state (111–114). Specifically, phosphorylation of ezrin at T567 promotes an open, active conformation that anchors the plasma membrane to the actin cytoskeleton, while dephosphorylation of T567 favors a closed, inactive conformation that weakens this linkage (111–114). Work from the Danuser lab has shown that dephosphorylation of membrane-associated ezrin releases the plasma membrane from actin to initiate cell protrusions, a mechanism that may also regulate PAP membrane dynamics and plasticity (115). Consistent with this idea, the Eroglu lab showed that modulating levels of ERM phosphorylation affects astrocyte morphological complexity (116).

Immunostain analysis of cocultures for p-ERMs revealed punctate and filopodial-like structures that were restricted to mature astrocytes (Fig. S1D, Fig. S5I; DIV7). In less mature astrocytes, p-ERM-positive structures often adopted a broader, fan-like morphology, consistent with a role in lamellipodial dynamics rather than synaptic ensheathment (Fig. S5I; DIV3). As astrocytes matured in coculture, the density of p-ERM-positive structures increased along astrocyte branches, and these structures were often found proximal to the postsynaptic marker PSD-95 (Fig. S5J-K). Thus, cocultured astrocytes form PAP structures, key features of astrocyte-synapse interactions.

Having established that PAPs form in coculture, we next asked whether astrocytic lysosome positioning is sensitive to synaptic activity. Strikingly, inhibiting glutamatergic signaling with CNQX and AP5 increased LEL motility in astrocyte branches (Fig. S6). Thus, LEL dynamics in cocultured astrocytes are coupled to glutamatergic signaling, suggesting that astrocytic lysosomes may respond to synaptic cues. To understand how astrocytic lysosomes might engage with synaptic compartments, we first mapped their spatial relationship to PAPs. We transduced cocultures with lentiviral LifeAct-mNG expressed under a truncated GFAP promoter to visualize F-actin within mature astrocyte branches. Similar to Fig. 4B, we observed the enrichment of F-actin in hot spots along the shaft of the astrocyte branch and in fine protrusions off the branch axis (Fig. 5A-Aii). Sites of F-actin concentration were also enriched for p-ERM (Fig. 5A-Ai) and the plasma membrane glutamate transporter GLAST (Fig. 5A, Aii), consistent with PAP structures described *in vivo*. Interestingly, LAMP1-positive LELs associated at foci enriched for actin, GLAST, and pERM, suggesting that LELs may be positioned to locally influence PAP structures (Fig. 5B; arrowheads).

Given the connections between lysosomal TRPML1 and actin, we investigated whether TRPML1 modulated p-ERM in astrocyte PAPs. Indeed, treatment with ML-SA1 (60 µM for 5 minutes) induced a rapid decrease in p-ERM area along astrocyte branches, relative to the DMSO solvent control, as measured by immunostain (Fig. 5C, D, Di). To test whether this effect reflected a change in phosphorylation versus total ERM, we turned to immunoblotting, since a reliable antibody against total ERM proteins for immunostaining was not available. Importantly, immunoblot analysis of monocultured astrocytes confirmed that ML-SA1 selectively reduced ERM phosphorylation without altering total ERM levels (Fig. 5G, H; Fig. S7A-D). Another TRPML1 agonist, MK6-83, elicited a similar trend of decreasing p-ERM levels relative to total ERM protein (Fig. S7E-H). Conversely, treatment with ML-SI3 elicited the opposite effect and rapidly increased p-ERM area along the astrocyte branch relative to the DMSO solvent control, as measured by immunostain (Fig. 5E, F, Fi). Immunoblot analysis confirmed no change in total ERM protein with ML-SI3 treatment (Fig. 5G, H). Interestingly, a time course analysis revealed that this increase in p-ERM area evoked by ML-SI3 was rapid and transient, peaking at 5 minutes followed by a recovery (Fig. S7I-K).

Combined, these data demonstrate that TRPML1 can rapidly modulate the phosphorylation of actin-membrane linkers enriched in PAPs; activation of TRPML1 dampens p-ERM levels and inhibition of TRPML1 increases p-ERM levels. Notably, these changes in p-ERM levels are transient, suggesting a dynamic, reversible process where TRPML1 may rapidly tune PAP membrane fluidity. Our data support a model in which arrested LELs serve site-specific functions that may influence PAP membrane dynamics at astrocyte-synapse interfaces (Fig. 6).

**Figure 6.**
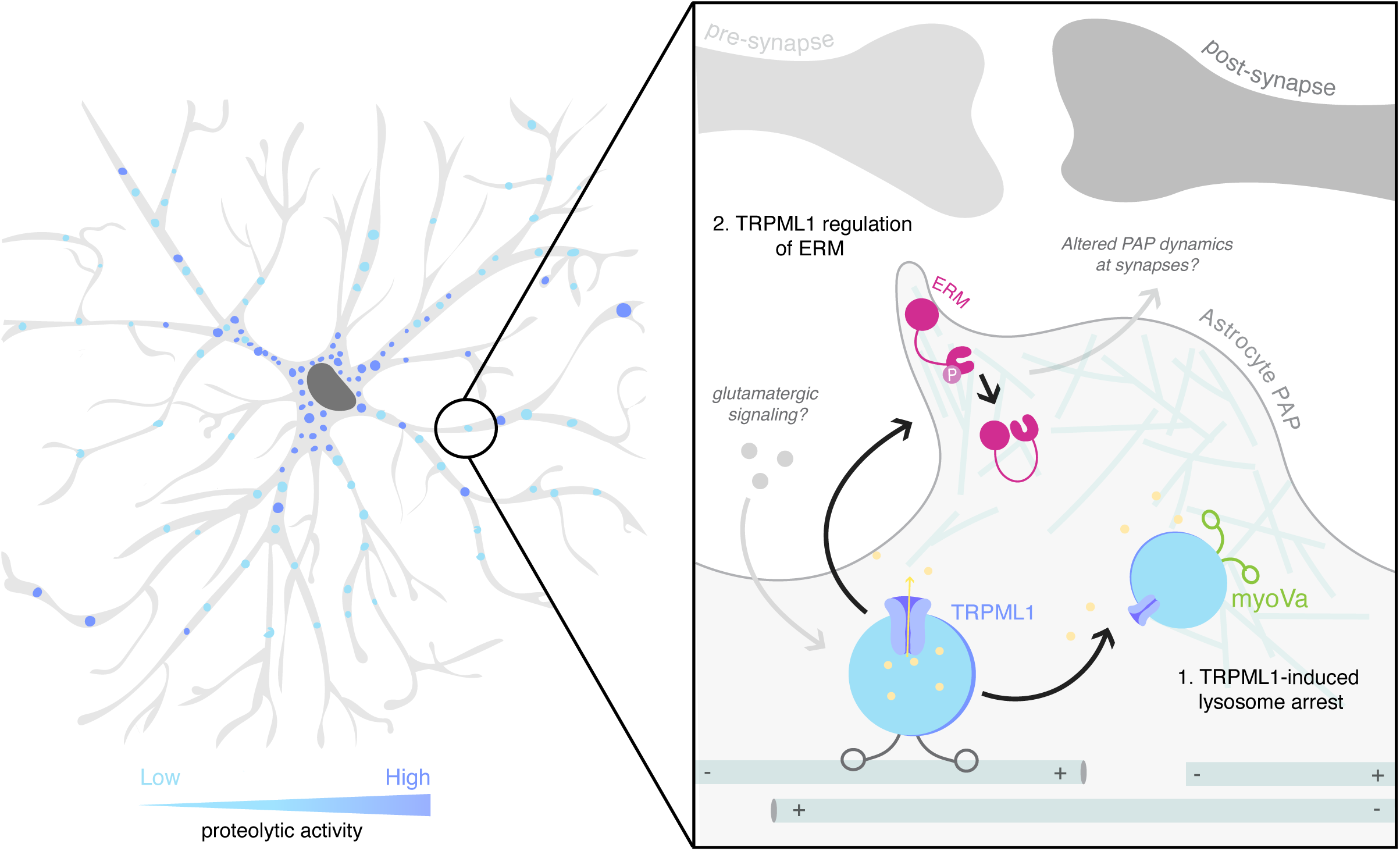
Model for TRPML1-dependent coordination of LEL positioning and regulation of the PAP cytoskeleton in astrocytic branches. LELs establish a spatial degradative gradient in astrocytic branches, with degradative activity concentrated in the soma and proximal branches. We have identified two key functions for lysosomal TRPML1. (1) TRPML1 activity promotes LEL motility arrest through a mechanism dependent on F-actin and myosin-Va. LEL tethering is coupled to glutamatergic signaling, suggesting that neuronal cues could influence LEL positioning within astrocytic branches. (2) TRPML1 signaling rapidly modulates ERM phosphorylation within PAPs. Changes in ERM phosphorylation may in turn influence PAP structural dynamics at the synapse.

## DISCUSSION

Our study identifies TRPML1 as a key regulator of LEL motility that, through engagement with the actin cytoskeleton, may position lysosomes to locally regulate actin-membrane linkers in PAPs. We show that LELs establish a gradient of degradative activity during branch maturation, with activity concentrated in the soma. Moreover, LELs within astrocyte branches undergo predominantly bidirectional movement that becomes progressively dampened as branches mature. TRPML1 is a driver of LEL immobilization in astrocyte branches. Mechanistically, TRPML1 activation promotes LEL arrest via myosin-Va-dependent tethering to the actin cytoskeleton, which may position LELs in proximity to PAPs. In parallel, TRPML1 activity regulates ERM protein phosphorylation in a rapid and transient manner, implicating lysosomal calcium signaling as an upstream regulator of the cytoskeletal machinery implicated in PAP membrane dynamics. Together, these findings suggest that TRPML1 acts as a dual regulator, controlling both LEL positioning and the activity of actin-associated membrane linkers, that may influence PAP structural plasticity.

TRPML1 likely coordinates both of these functions by mediating lysosomal calcium efflux, consistent with an emerging role for lysosomes shaping both local and global calcium events (117). Astrocytes utilize complex calcium fluctuations over a wide spatiotemporal scale; from discrete microdomain events in PAPs (∼msec), to waves of calcium that can engage the soma and surge throughout the cell (∼sec) (118). Astrocyte calcium events are intricately coupled to neuronal activity, and alterations in astrocytic calcium have functional consequences on circuits and behavior (119). For example, the structural remodeling of PAPs is calcium dependent (13, 18, 23). Indeed, chelating calcium reduces PAP motility, while inducing calcium transients within PAPs is sufficient to trigger PAP motility and subsequent coverage of associated spines (13). The endoplasmic reticulum (ER) has been proposed as the primary calcium source for this response (18, 120, 121). However, astrocyte-specific knockout of IP3R2, the predominant ER calcium release channel in astrocytes, significantly reduces somatic calcium fluctuations, but largely preserves calcium waves and microdomain transients in astrocyte processes, suggesting alternative calcium sources (119, 122, 123). Notably, lysosomal calcium concentrations are comparable to those in the ER (80) but have received very little attention in astrocytes. Work in *Drosophila* suggests a conserved role for the TRPML1 homolog TrpML in mediating calcium microdomain events in astrocytic processes evoked by neurotransmitters (124). Our data further support a role for LELs as an additional, spatially discrete calcium source distributed throughout astrocyte branches.

Thus, we propose that LELs may serve as mobile calcium carriers in astrocyte branches, with TRPML1 positioning these stores to support perisynaptic functions such as calcium-dependent PAP motility. Consistent with this idea, neurons in Huntington’s disease models exhibit both impaired LTP and depletion of LELs from dendritic compartments (125). Optogenetic repositioning of LELs into dendrites restores LTP via local lysosomal calcium signaling (125). Moreover, TRPML1 activity can enhance neuronal excitability and partially rescue synaptic defects in a mouse model of the lysosomal storage disorder Juvenile Neuronal Ceroid Lipofuscinosis (JNCL or Batten Disease) (126). Thus, accumulating evidence supports a role for lysosomal calcium signaling in synaptic support. In our study, the tethering of LELs through TRPML1 may position a locally releasable calcium source at sites proximal to PAPs. However, future work will need to resolve if TRPML1-dependent changes in ERM phosphorylation within PAPs are a direct consequence of local LEL positioning or the result of a branch-wide calcium signaling event.

Whether TRPML1 activity is itself sensitive to neuronal activity is an important question. Consistent with this possibility, we observed that glutamatergic signaling restricted LEL motility in astrocyte branches, in a manner similar to pharmacological TRPML1 activation, suggesting that astrocytic lysosomal trafficking is responsive to neuronal signals. This coupling to synaptic activity aligns LEL behavior in astrocyte branches with neuronal dendrites. In both compartments, LEL motility is dampened by glutamatergic signaling (1, 2), and in dendrites this arrest involves myosin-Va tethering to actin hotspots along the dendritic shaft, proximal to spines (84). Since myosin-Va activity is regulated by calcium and calmodulin (98–100), TRPML1 signaling in astrocytes could provide the local upstream calcium signal needed to coordinate LEL tethering along branches. This activity-dependent tethering could position astrocytic LELs to locally regulate PAP structural plasticity at the astrocyte-synapse interface.

Similar to our observations in astrocyte branches, dendrites exhibit a proximal-distal gradient of degradative activity in mature processes, and LELs in dendrites also undergo primarily short-range bidirectional motility (37, 127). This shared motility behavior between astrocyte branches and neuronal dendrites likely reflects a common underlying mechanism: a microtubule network of mixed polarity. As dendrites mature, microtubule polarity shifts from unidirectional to mixed (in the proximal 2/3’s of the dendrite) (128–132). Similarly, as astrocyte branches mature, we found that microtubule polarity becomes more mixed (∼2/3 plus-end-out), a transition that corresponded with decreased LEL motility and an increased fraction of stationary LELs. Notably, the mixed microtubule polarity we observed in mature astrocyte branches contrasts with the predominantly uniform polarity reported in monocultured astrocytes (133), suggesting that maturation state and cellular environment may shape cytoskeletal organization. Indeed, neurons provide instructive cues that influence the development and maturation of local astrocytes (134–137), and their presence in our coculture system may contribute to this cytoskeletal reorganization. Interestingly, TRPML1 also regulates lysosomal transport in neurons, though its effects vary by compartment; TRPML1 activation attenuates LEL motility in axons (55) but stimulates LEL motility in dendrites (56, 57). Future work will need to resolve how the same molecular signal can be interpreted differently depending on local context.

We find that activation of TRPML1 rapidly promotes both LEL immobilization and ERM dephosphorylation. This tethering may position LELs to transiently regulate the activity of ERM proteins which may trigger morphological changes in PAPs. Ezrin levels and the degree of ezrin phosphorylation are critical determinants of astrocyte morphology and structural plasticity *in vivo* (16, 116, 138–140). Both hyper- and hypo-phosphorylation of ERM proteins are associated with reduced astrocyte territory volume and morphological complexity, phenotypes rescued by phospho-dead or phospho-mimetic ezrin substitutions at T567, respectively (116). Flexibility in the phosphorylation state of ezrin is therefore likely required for the dynamic extension and retraction of PAPs. Consistent with this idea, acute local dephosphorylation of ezrin T567 at the plasma membrane is sufficient to initiate membrane protrusion in cultured mammalian cells, with actin polymerization providing the mechanical force to advance the membrane (115). Thus, release of actin-membrane connections enables membrane fluidity or deformation. Under conditions requiring PAP membrane stabilization, the balance may shift towards ERM phosphorylation, promoting actin-membrane linkage (21). Similar patterns have been reported in the filopodia of neuronal growth cones, where transient elevations in p-ERM slow neurite outgrowth and reduced p-ERM promote outgrowth (141). We find that the effects of modulating TRPML1 on p-ERM levels are rapid (within 5 minutes) and transient, consistent with the temporal profile of PAP dynamics. TRPML1-driven changes in p-ERM may transiently decrease membrane-actin coupling in astrocyte branches, rendering the membrane more deformable. Whether this permissive state results in filopodial expansion or retraction will require future studies that monitor membrane dynamics in tandem with the underlying actin cytoskeleton.

Our observation that TRPML1 bidirectionally regulates ERM phosphorylation raises the possibility that lysosomal calcium contributes to rapid cytoskeletal remodeling underlying PAP plasticity. Future work will need to explore the signaling pathways that connect TRPML1-mediated calcium efflux to changes in ERM phosphorylation and potential downstream effects on the actin cytoskeleton. Lysosomal calcium release via TRPML1 can activate the phosphatase calcineurin (142), and calcineurin can affect actin dynamics via downstream targets such as cofilin, an actin-severing and depolymerization factor which has been implicated in PAP remodeling (15, 143). However, a direct connection between these molecular players has not yet been established in astrocytes. The kinases and phosphatases that regulate ERM activity have been characterized in non-neural cell types (111), but are largely unknown in astrocytes. However, recent evidence links the Parkinson’s disease-associated Leucine-Rich Repeat Kinase 2 (LRRK2) gain-of-function mutation (G2019S) with ERM hyperphosphorylation, altered astrocyte morphology, and synaptic deficits, underscoring ERM dysregulation as a pathological vulnerability in astrocytes (116).

Links between TRPML1 and the actin cytoskeleton were largely unanticipated, yet precedent for such coupling has been established in the fields of immunology and cancer biology (106–108, 144, 145). Several studies have shown that TRPML1 is required for immune cell migration by controlling actomyosin contractility at the cell rear (106–108). Mechanistically, TRPML1 drives the redistribution of lysosomes to the rear of the cell and reorganizes the actin cytoskeleton (106–108). Lysosomal calcium release locally activates myosin-II, thereby generating the actomyosin contractile forces that propel cell movement (106–108). Consistent with these findings, TRPML1 also promotes the migration of cancer cells by regulating actin polymerization and the trafficking of cell adhesion molecules (144). In highly aggressive cancer models, TRPML1 promotes cell migration and invasive potential by stimulating actin polymerization and peripheral lysosome positioning, triggering lysosomal secretion of proteases which may remodel the extracellular matrix (145). Thus, roles for TRPML1 in coordinating changes in the actin cytoskeleton with membrane dynamics is emerging as a conserved mechanism across various cell types. While mature astrocytes are largely not migratory, TRPML1-dependent regulation of the actin cytoskeleton may have been harnessed for other forms of structural remodeling, such as the morphological plasticity of PAPs.

LEL immobilization might also function to locally concentrate degradative activity in response to synaptic signals. In dendrites, synaptic activity stimulates lysosomal degradation, remodeling the local synaptic proteome (1–4, 57, 146). Branch-localized LELs in astrocytes likely also serve an analogous function in local degradation. Additionally, lysosomes are equipped with amino acid exporters that release proteolytic products into the cytoplasm, thereby converting degradation products to biosynthetic fuel for local protein synthesis (80). LELs can also associate with mRNAs, further contributing to the molecular machinery that drives local protein translation (147, 148). A diversity of transcripts (e.g., metabolic, cytoskeletal, and synapse-associated) are locally translated within PAPs (149–151). Thus, LELs in astrocyte branches might support local synthesis of PAP proteins. In this way, branch-localized LELs may coordinate both sides of the metabolic equation, balancing local degradation with protein synthesis, in a process of creation by destruction.

TRPML1 may also tether LELs along the branch, staging these organelles at sites poised for lysosomal secretion. We observed that branch-localized LELs are less degradative than those in the soma, consistent with observations in non-polarized cells where peripheral LELs are less degradative and potentially more exocytic (152–154). Indeed, activating TRPML1 stimulates lysosomal exocytosis in monocultured astrocytes (61). TRPML1-mediated lysosomal secretion may serve several functions. First, it might provide additional membrane to support expanding PAP structures (155). Alternatively, LELs contain high concentrations of ATP that can be released extracellularly in response to glutamatergic stimulation (31, 32, 61). Lysosomal ATP has been proposed as a mechanism by which astrocytes modulate purinergic signaling at synapses (30, 156–160), although direct evidence of lysosomal secretion in this context remains limited. Interestingly, astrocyte-specific knockout of TRPML1 in the medial prefrontal cortex reduced ATP release *in vivo* and was sufficient to induce depressive-like behaviors in mice (26), supporting a role for astrocytic TRPML1 in lysosomal ATP secretion that modulates synaptic function. The peripheral positioning of LELs in astrocyte branches is likely crucial for this ATP release. For example, iPSC-derived astrocytes of Alexander’s disease accumulate lysosomes in the perinuclear region (due to disrupted intermediate filaments) and exhibit impaired ATP release (33).

In sum, TRPML1 may position LELs along astrocyte branches to support multiple functions including local calcium signaling, degradation, and secretion, that collectively influence astrocyte morphology and synaptic interactions. Key next steps will be to assess PAP motility and synaptic coverage, particularly *in vivo,* to resolve the structural and functional consequences of transient changes in TRPML1 signaling.

Our study establishes foundational pathways by which TRPML1 regulates lysosomal trafficking and function in astrocytes, with potential implications for astrocyte-synapse contacts and synaptic function. An important next step will be to determine how these pathways are disrupted in Mucolipidosis type IV (MLIV), where loss-of-function mutations in TRPML1 impair lysosomal function. In *MCOLN1*-/- mice, gliosis of astrocytes precedes excitotoxicity or neurodegeneration, implying an early glial insult in the disease (52). In a *Drosophila* model of MLIV, expression of wild-type TrpML in glia is sufficient to rescue synaptic and behavioral deficits (161). Combined, these models suggest a strong glial involvement in MLIV progression, with dysfunction in astrocytic lysosomes central to pathology.

Studies of the aging mouse brain also implicate defective astrocytic lysosomes in driving synaptic pathology in neurodegenerative diseases (25). An age-dependent subpopulation of astrocytes was found in the hippocampus that accumulates swollen autophagic-lysosomal compartments within branches, coinciding with disrupted synapse formation near affected astrocyte territories (25). Models of Alzheimer’s disease showed elevated levels of these astrocyte populations (25). Thus, lysosomal dysfunction in astrocytes is emerging as a key contributor to neurological disease and may represent a new avenue for therapeutic intervention. Our work provides a mechanistic framework for beginning to understand how lysosomal dysfunction in astrocytes may drive synaptic pathology in disease.

## AUTHOR CONTRIBUTIONS

Conceptualization, M.L.S. and S.M.; Methodology, M.L.S., M.M., S.M.; Validation, M.L.S., S.M.; Investigation and Formal Analysis, M.L.S., M.L.F., S.C., D.S., Y.W., J.B.; Resources, S.M.; Data Curation, M.L.S., M.L.F., Y.W., J.B., M.M, S.M.; Writing - Original Draft, M.L.S., S.M.; Writing - Review and Editing, M.L.S., S.M.; Visualization, M.L.S., Y.W., J.B., M.M., S.M.; Supervision, Project administration, and Funding Acquisition, S.M..

## ACKNOWLEDGEMENTS

This work was supported by NIH grants R01NS110716 (to S.M.) and F31NS132453 (to M.L.S.). We thank Dr. Mary Putt (Biostatistics and Bioinformatics Core at the Intellectual and Developmental Disabilities Research Center at the Children’s Hospital of Philadelphia) and Dr. Edward Lee (Perelman School of Medicine at the University of Pennsylvania) for advice with statistical analyses. We thank Dr. Kelly Jordan-Sciutto, Dr. Lindsay Festa, and Dr. Judith Grinspan for insightful discussions and helpful feedback on the study. We thank Dr. James Shorter for critical feedback on the manuscript.

---

**Figure S1.**
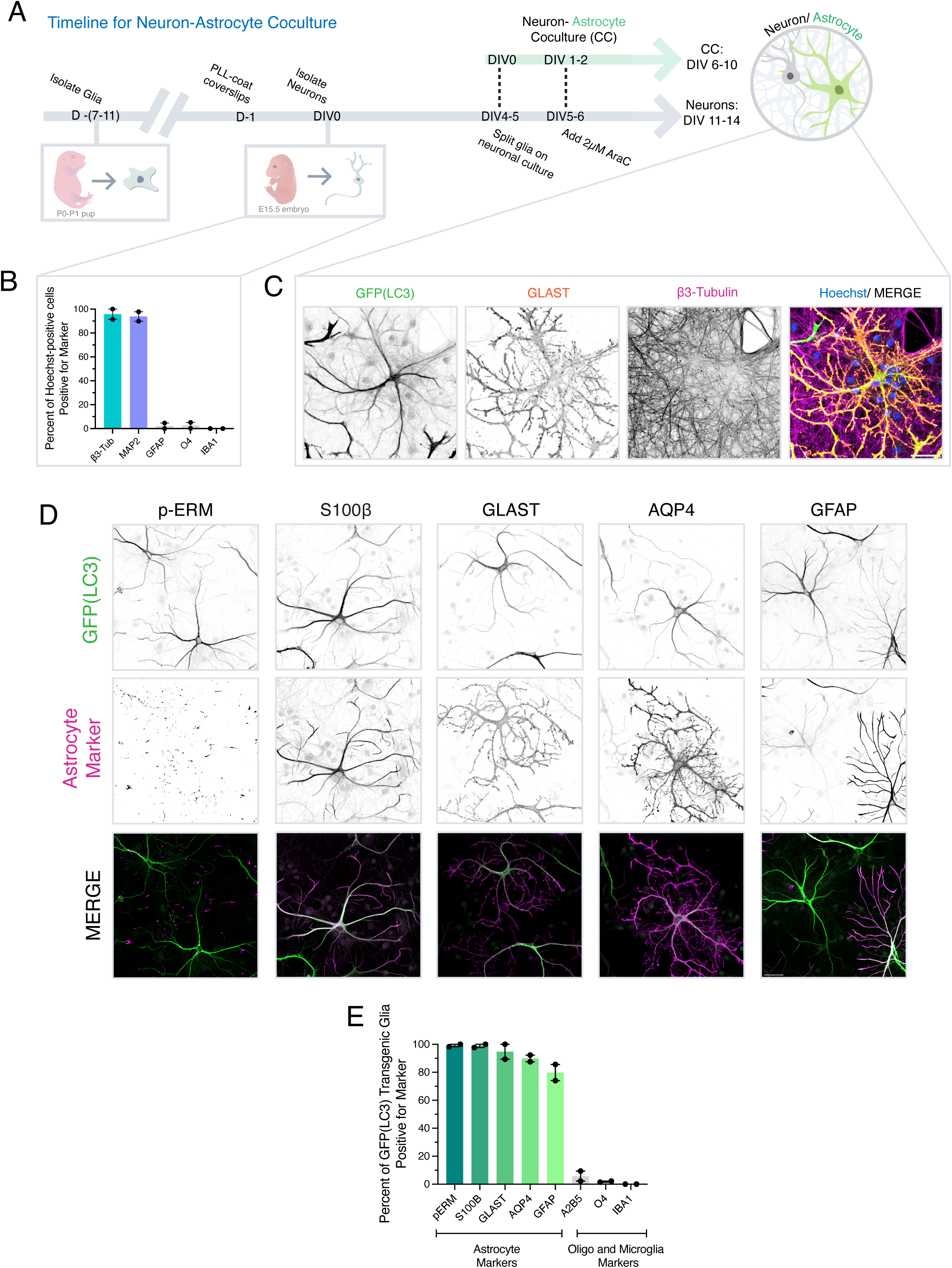
A cortical murine neuron-astrocyte coculture system to study stellate astrocytes in vitro. **(A)** Schematic of the protocol for neuron-astrocyte coculture. Cortical glia enriched for astrocytes are isolated from postnatal day 0-1 (P0-1) pups. To distinguish astrocytes by live-cell imaging, cortical glia are often isolated from GFP-LC3 transgenic mice (for subsequent coculture with non-transgenic neurons). The day prior to neuron isolation, acid-washed coverslips are coated with poly-L-lysine. Cortical neurons are isolated at embryonic day 15.5, plated onto PLL-coated coverslips, and maintained as a monoculture for 4-5 days in vitro (DIV), at which point cortical glia are plated onto the neuronal meshwork (designated as DIV0 of coculture). One to two days after addition of glia, cytosine arabinoside (AraC; 2 µM) is added to the coculture to restrict astrocyte proliferation. Cocultures are maintained for a total of 6-10 days (neurons are a total of DIV11-14). **(B)** Quantification of the purity of the neuron monoculture. Shown is the percentage of nuclei in non-transgenic cortical neuronal monocultures at DIV8 that are co-positive for markers of neurons (β3-Tubulin to label neuronal microtubules, MAP2 to label dendrites), astrocytes (GFAP), oligodendrocytes (O4), or microglia (Iba1). A total of 4,198 cells were counted across 2 biological replicates. **(C)** Representative maximum-intensity projections of z-stacks of the neuron-astrocyte coculture at DIV7 of coculture. Non-transgenic neurons (total age of DIV11) were cocultured with GFP-LC3 transgenic glia and immunostained for β3-Tubulin (neuron marker), GFP (to visualize LC3-expressing glia), and GLAST to confirm astrocyte identity. Scale bar, 20 µm. **(D-E)** Quantification of the identity of GFP-LC3-expressing glia in coculture with neurons. **(D)** Representative maximum-intensity projections of z-stacks of the neuron-astrocyte coculture at DIV7 of coculture. Cocultures of non-transgenic neurons and GFP-LC3 transgenic glia were immunostained for GFP (to label the GFP-LC3 transgene only present in glia) and a panel of astrocyte markers: p-ERM (Ezrin [Thr567]/Radixin [Thr564]/Moesin [Thr558]), S100β (S100 calcium-binding protein B), GLAST (excitatory amino acid transporter 1, EAAT1), AQP4 (aquaporin-4), and GFAP (glial fibrillary acidic protein). Scale bar, 40 µm. **(E)** Percentage of GFP-LC3 transgenic glia (cocultured for DIV7) co-positive for markers of astrocytes (p-ERM, S100β, GLAST, AQP4, GFAP), oligodendrocyte lineages (A2B5, O4), or microglia (Iba1). A total of 903 cells were counted across 2 biological replicates.

**Figure S2.**
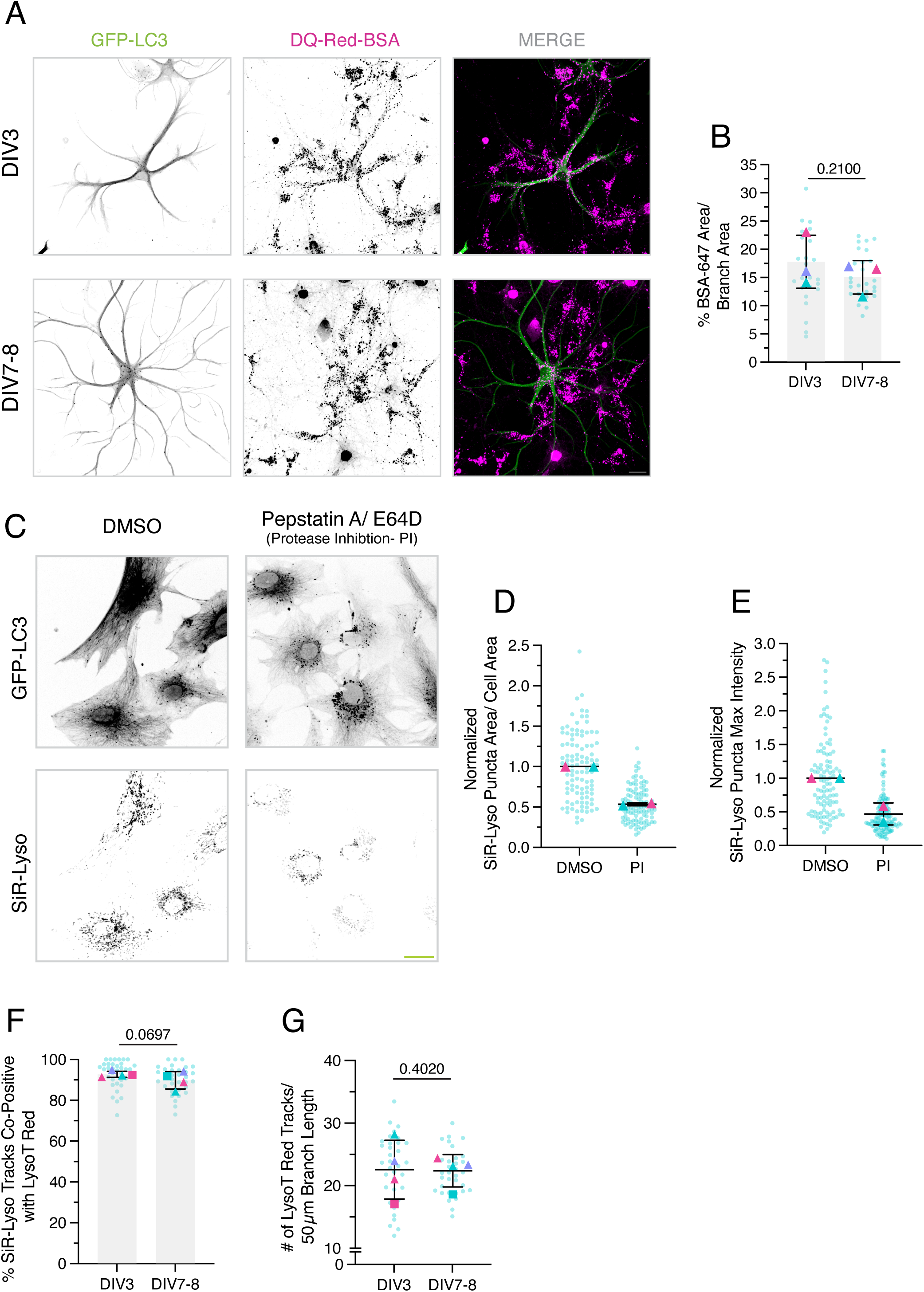
Validation of endolysosomal markers across astrocyte branch maturation. **(A)** Uncropped single channel and merge of images shown in Fig. 1A to visualize LEL signal in neighboring neurons; scale bar 20 µm. **(B)** The graph in B corresponds to the experiment and primary data in Figure 1A, C-D. Quantification of the total area occupied by BSA-647-positive puncta normalized to branch area; GFP-LC3 was used as a space-fill to delineate the branch boundary. Horizontal bars represent the mean of biological replicates ± SD; shown are p-values from a LME model; N=25-26 branches (one branch per astrocyte) across 3 biological replicates. **(C-E) (C)** Maximum-intensity projections of live-cell z-stacks from monoculture GFP-LC3 transgenic astrocytes at DIV6 labeled with SiR-Lysosome. Astrocytes were treated for 3 hours with DMSO (solvent control) or a protease inhibitor cocktail (PI; 30 µM Pepstatin-A and 30 µM E64D). Images within each channel are displayed with identical minimum and maximum intensity settings across treatment conditions. Scale bar, 40 µm. **(D)** Corresponding quantification of the total area occupied by SiR-Lysosome-positive puncta normalized to cell area. **(E)** SiR-Lysosome puncta maximum intensity; values are normalized to the mean of the DMSO-treated group within each replicate. **(D-E)** N=40-65 astrocytes from 2 biological replicates. **(F-G)** Graphs in F-G are paired with the analysis in Figure 1E-G, and correspond to the experiment and primary data for the DMSO controls at each time point derived from the dataset in Figure S4A-D. **(F)** The density of LysoTracker Red tracks normalized to a 50 µm branch segment of coculture astrocytes; the GFP-LC3 transgene was used as a space-fill to delineate the branch boundary. **(G)** Percentage of SiR-Lysosome tracks that are co-positive for LysoTracker Red. **(F-G)** Horizontal bars represent the mean of biological replicates ± SD; shown are p-values from a LME model; N=33-34 branches (one branch per astrocyte) across 4 biological replicates.

**Figure S3.**
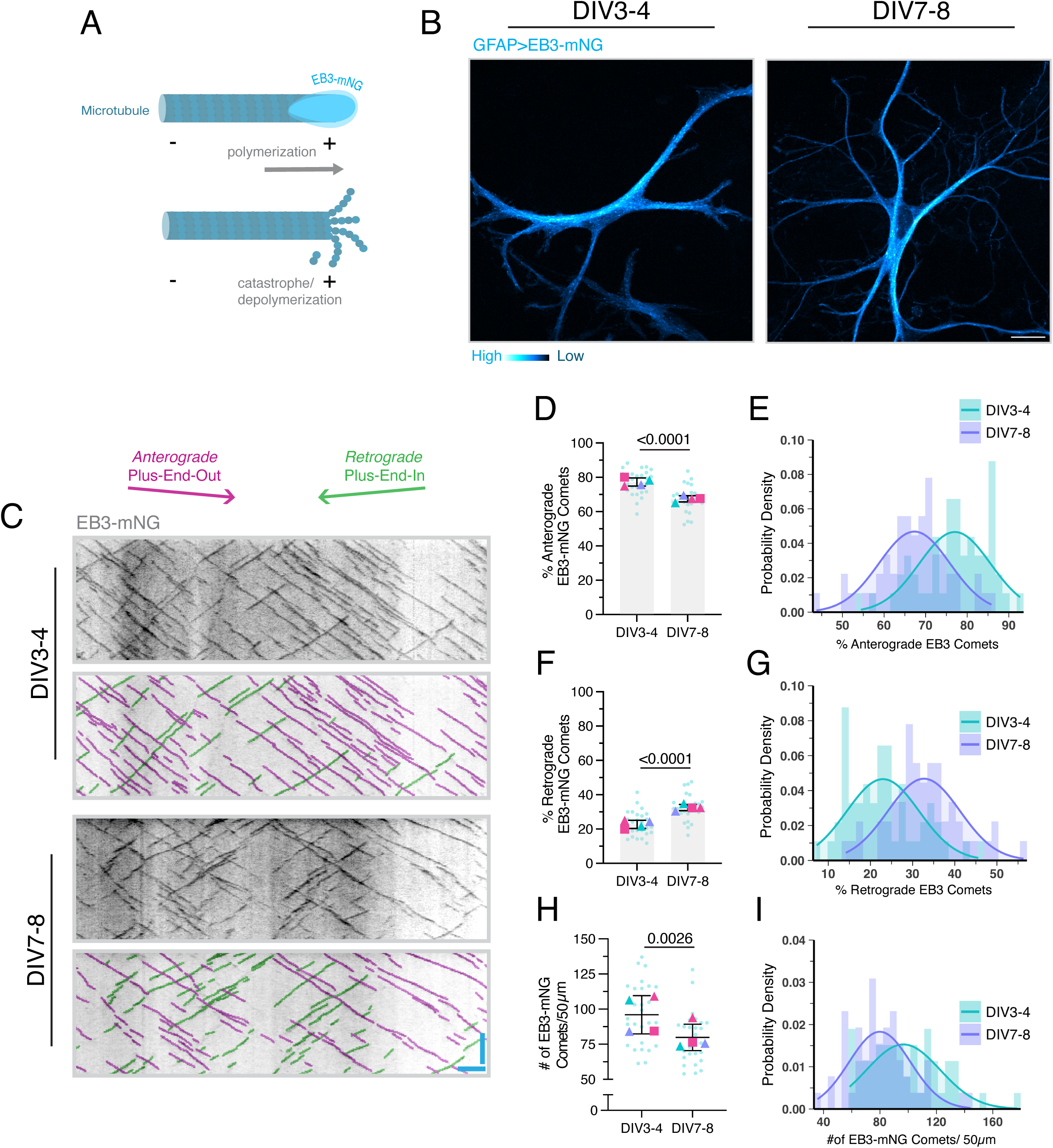
Microtubule polarity becomes more mixed as astrocyte branches develop in coculture with neurons. **(A)** Schematic of a dynamic microtubule. To track the directionality of polymerizing microtubules in astrocyte branches, we transduced non-transgenic cocultures with EB3-mNeonGreen (mNG) under a truncated GFAP promoter. EB3 (End-binding protein 3) specifically binds the GTP-tubulin-rich cap at the growing plus ends of microtubules. EB3 rapidly dissociates upon GTP hydrolysis during catastrophe/depolymerization. Thus, EB3-NG produces a comet-like fluorescent signal that tracks the plus-end of polymerizing microtubules. **(B)** Representative single-plane images of non-transgenic astrocytes expressing lentiviral EB3-mNeonGreen. Scale bar, 20 µm. **(C-I) (C)** Representative kymographs of EB3-mNeonGreen dynamics along a primary-to-secondary branch trajectory in astrocytes at coculture DIV3-4 versus DIV7-8. At each time point, top panels are raw kymographs; EB3 comets that were manually tracked (anterograde comets shown in magenta and retrograde comets shown in green) are overlaid onto the respective kymographs and shown in the bottom panels. All quantification is derived from the kymograph analysis. Horizontal bar, 5 µm; vertical bar, 30 sec. **(D)** Percentage of anterograde EB3-mNeonGreen tracks. Horizontal bars represent the mean of biological replicates ± SD; shown are p-values from a LME model; N=27 astrocytes across 4 biological replicates; each astrocyte value represents the mean of two branches. **(E)** Corresponding probability density histogram for data in D. Solid curves show kernel density estimates. N=53-54 individual branches (not averaged per astrocyte) across 4 biological replicates. **(F)** Percentage of retrograde EB3-mNeonGreen tracks. Horizontal bars represent the mean of biological replicates ± SD; shown are p-values from a LME model; N=27 astrocytes across 4 biological replicates; each astrocyte value represents the mean of two branches. **(G)** Corresponding probability density histogram with kernel density estimates for data in F. N=53-54 individual branches (not averaged per astrocyte) across 4 biological replicates. **(H)** Early electron microscopy studies reported that nascent astrocyte processes are densely populated with microtubules (162, 163). As astrocytes mature, however, these processes become enriched in intermediate filaments and comparatively depleted of microtubules (162, 163). Similarly, we observed a decrease in EB3 tracks in mature astrocyte branches. H shows the density of EB3-mNeonGreen tracks normalized to a 50 µm branch segment. Horizontal bars represent the mean of biological replicates ± SD; shown are p-values from a LME model; N=27 astrocytes across 4 biological replicates; each astrocyte value represents the mean of two branches. **(I)** Corresponding probability density histogram for data in H. N=53-54 individual branches (not averaged per astrocyte) across 4 biological replicates.

**Figure S4.**
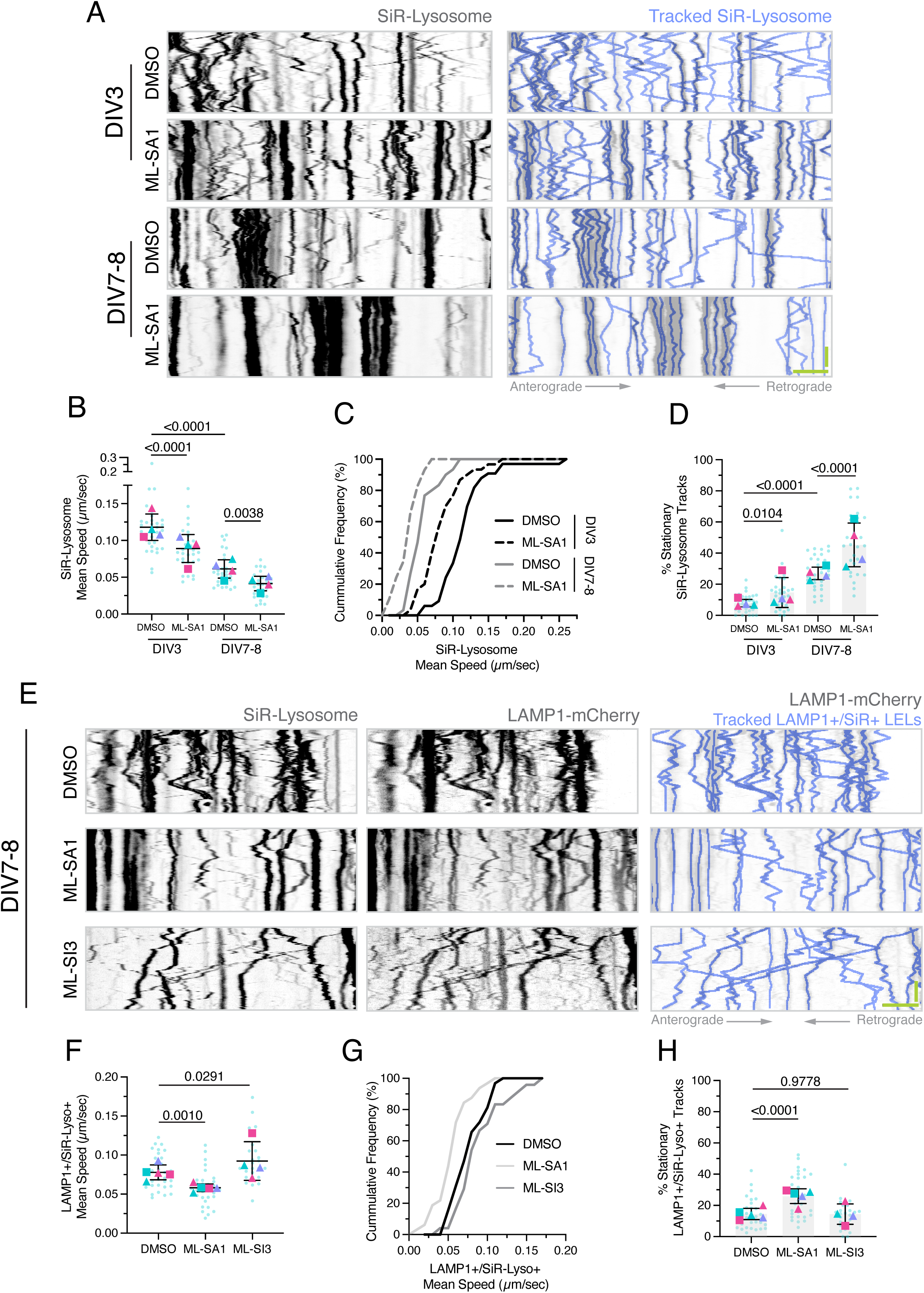
Lower concentrations of TRPML1 pharmacological modulators are sufficient to alter LEL motility across different stages of astrocyte branch maturation. (A-D) **(A)** Representative kymographs of SiR-Lysosome motility along astrocytic branches at DIV3 versus DIV7-8 of coculture; the GFP-LC3 transgene was used as a space-fill to delineate the branch boundary. Cocultures were treated for 2 hours with DMSO (solvent control) or ML-SA1 (20 µM). SiR-Lysosome puncta that were manually tracked (shown in periwinkle) are overlaid onto their respective kymographs. Horizontal bar, 5 µm; vertical bar, 30 sec. All quantification is derived from kymograph analysis: **(B)** mean SiR-Lysosome track speed (µm/sec), **(C)** cumulative frequency of SiR-Lysosome mean speed (derived from the technical replicates in B), and **(D)** percentage of stationary (or immobile) tracks**. (B, D)** Horizontal bars represent the mean of biological replicates ± SD; shown are p-values from a LME model with Holm’s correction for multiple comparisons; N=29-33 branches (one branch per astrocyte) across 4 biological replicates. Data from the DMSO condition in Figure S4A-D are also analyzed in Figure 1E-G and I-K. **(E-H) (E)** Representative kymographs of SiR-Lysosome and lentiviral LAMP1-mCherry (expressed under a truncated GFAP promoter) motility along astrocytic branches cocultured for DIV7-8; lentiviral eGFP (co-expressed under a truncated GFAP promoter) was used as a space-fill to delineate the branch boundary. Cocultures were treated for 2 hours with DMSO (solvent control), ML-SA1 (20 µM), or ML-SI3 (10 µM). SiR-Lysosome puncta that were manually tracked (shown in periwinkle) are overlaid onto their respective kymographs. Horizontal bar, 5 µm; vertical bar, 30 sec. All quantification is derived from kymograph analysis: **(F)** mean SiR-Lysosome track speed (µm/sec), **(G)** cumulative frequency of SiR-Lysosome mean speed (derived from the technical replicates in F), and **(H)** percentage of stationary (or immobile) tracks**. (F, H)** Horizontal bars represent the mean of biological replicates ± SD; shown are p-values from a LME model; N=24-32 branches (one branch per astrocyte) across 4-5 biological replicates. Data from the DMSO condition in Figure S4E-H are also analyzed in Figure 1L-N.

**Figure S5.**
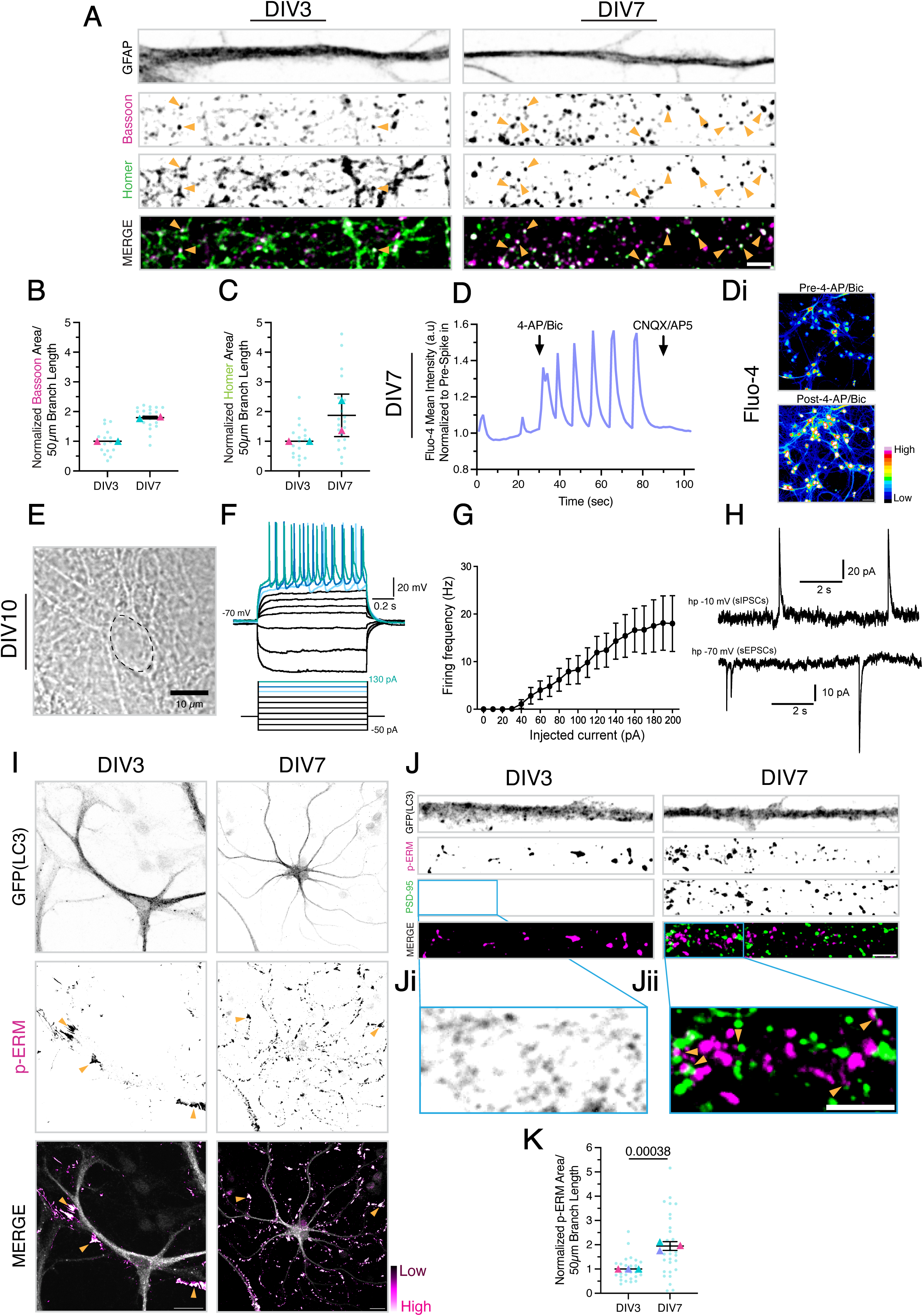
Neurons form mature synapses and astrocytes form elaborate PAP structures in coculture. (A-D) **(A)** Straightened branches from a single plane of non-transgenic cocultures at DIV3 versus DIV7; astrocytes were identified by immunostain for GFAP. Cocultures were co-stained for Bassoon to label presynaptic compartments and Homer to label postsynaptic compartments. Images within each channel are displayed with identical minimum and maximum intensity settings across DIV. Filled orange arrowheads denote bona fide synapses that are co-positive for both pre- and postsynaptic markers, consistent with synaptic contacts adjacent to the astrocyte branch. Scale bar, 5 µm. Corresponding quantification of total area occupied by **(B)** Bassoon puncta or **(C)** Homer puncta. Data are normalized to a 50 µm branch segment and expressed as a fold change relative to the mean of the DIV3 corresponding biological replicate. **(B-C)** Horizontal bars represent the mean of biological replicates ± SD; N=17-20 branches (one branch per astrocyte) across 2 biological replicates. **(D-Di) (D)** Non-transgenic coculture at DIV7 (neurons are a total of DIV11) loaded with Fluo-4 AM and treated with 50 µM 4-AP and 50 µM bicuculline added via on-scope delivery between 29-30 seconds of live cell imaging, followed by on-scope addition of 50 µM CNQX and 50 µM AP5 added between 89-90 seconds. Trace represents the fluorescence mean intensity measured of the neuronal network, normalized to the average value for mean intensity of frames prior to spike in (0-29 seconds). **(Di)** Images corresponding to frames 1 and 39 from the trace in (D). **(E-H) (E)** Brightfield image of a neuron at DIV10 of coculture (total neuron age of DIV14) used for patch-clamp recording; soma is outlined by a black dashed contour. **(F)** Fired action potentials upon current injection (−50 to +130 pA in 20 pA steps). Under current-clamp mode, the baseline membrane potential was held at −70 mV. **(G)** Firing rates plotted against current injection (n=11 neurons at DIV10 of coculture). **(H)** All neurons exhibit sIPSCs (upper row: 7 out of 7 neurons recorded; holding potential = −10 mV) and sEPSCs (lower row: 14 out of 14 neurons recorded; holding potential = −70 mV) under voltage-clamp mode. **(I-K) (I)** Representative maximum-intensity projections of z-stacks of cocultured non-transgenic neurons and GFP-LC3 transgenic astrocytes at DIV3 versus DIV7, immunostained for GFP(LC3) and p-ERM. Orange arrowheads denote fan-like lamellate structures. Scale bar, 20 µm. **(J)** Straightened branches from maximum-intensity projections of z-stacks of cocultured non-transgenic neurons and GFP-LC3 transgenic astrocytes at DIV3 versus DIV7. Cocultures were immunostained for GFP (LC3; labels astrocytes), p-ERM, and the postsynaptic marker PSD-95. Images within each channel are displayed with identical minimum and maximum intensity settings across time points. Scale bar, 5 µm. **(Ji)** Higher-magnification view of PSD-95 with display settings adjusted independently to visualize low-level signal at DIV3 of coculture. **(Jii)** Higher-magnification view of p-ERM and PSD-95 highlighting regions of close apposition between the two signals (filled orange arrowheads). Scale bar, 5 µm. **(K)** Corresponding quantification of total area occupied by p-ERM-positive structures per unit branch length and expressed as a fold change relative to the mean of the DIV3 corresponding biological replicate. Horizontal bars represent the mean of biological replicates ± SD; shown are p-values from an LME model; N=29 branches (one branch per astrocyte) across 3 biological replicates.

**Figure S6.**
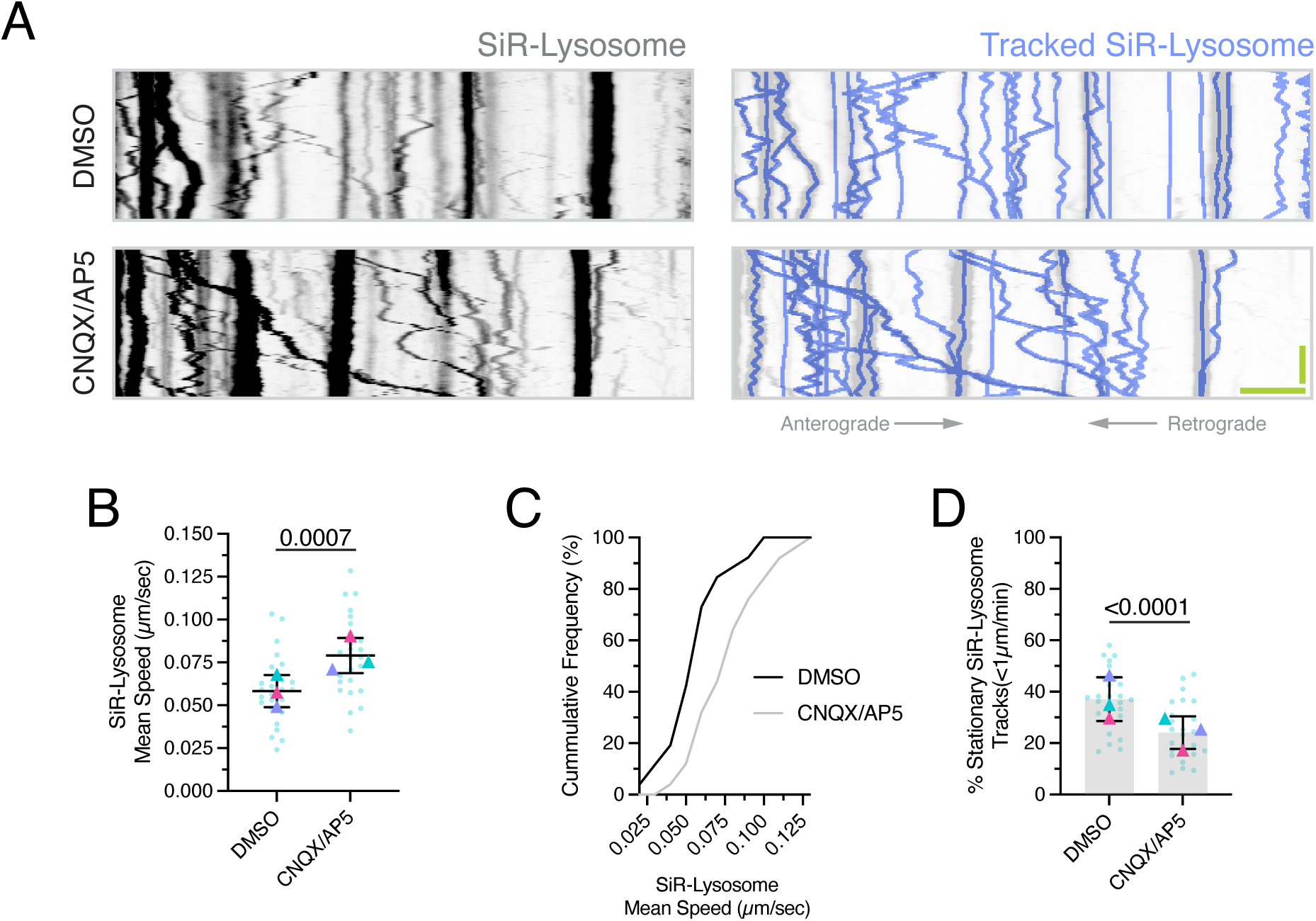
Glutamatergic signaling dampens LEL motility in astrocytic branches. (A-D) **(A)** Kymographs of SiR-Lysosome motility along astrocytic branches at DIV7 of coculture; the GFP-LC3 transgene was used as a space-fill to delineate branch boundaries of astrocytes in coculture with non-transgenic neurons. Cocultures were treated for 30 minutes with DMSO (solvent control) or CNQX/AP5 (50 µM/50µM). SiR-Lysosome puncta that were manually tracked (shown in periwinkle) are overlaid onto their respective kymographs. Horizontal bar, 5 µm; vertical bar, 30 sec. All quantification is derived from kymograph analysis: **(B)** mean SiR-Lysosome track speed (µm/sec), **(C)** cumulative frequency of SiR-Lysosome mean speed (derived from the technical replicates in B), **(D)** percentage of stationary (or immobile) tracks. **(B, D)** Horizontal bars represent the mean of biological replicates ± SD; shown are p-values from an LME model; N=25-26 branches (one branch per astrocyte) across 3 biological replicates.

**Figure S7.**
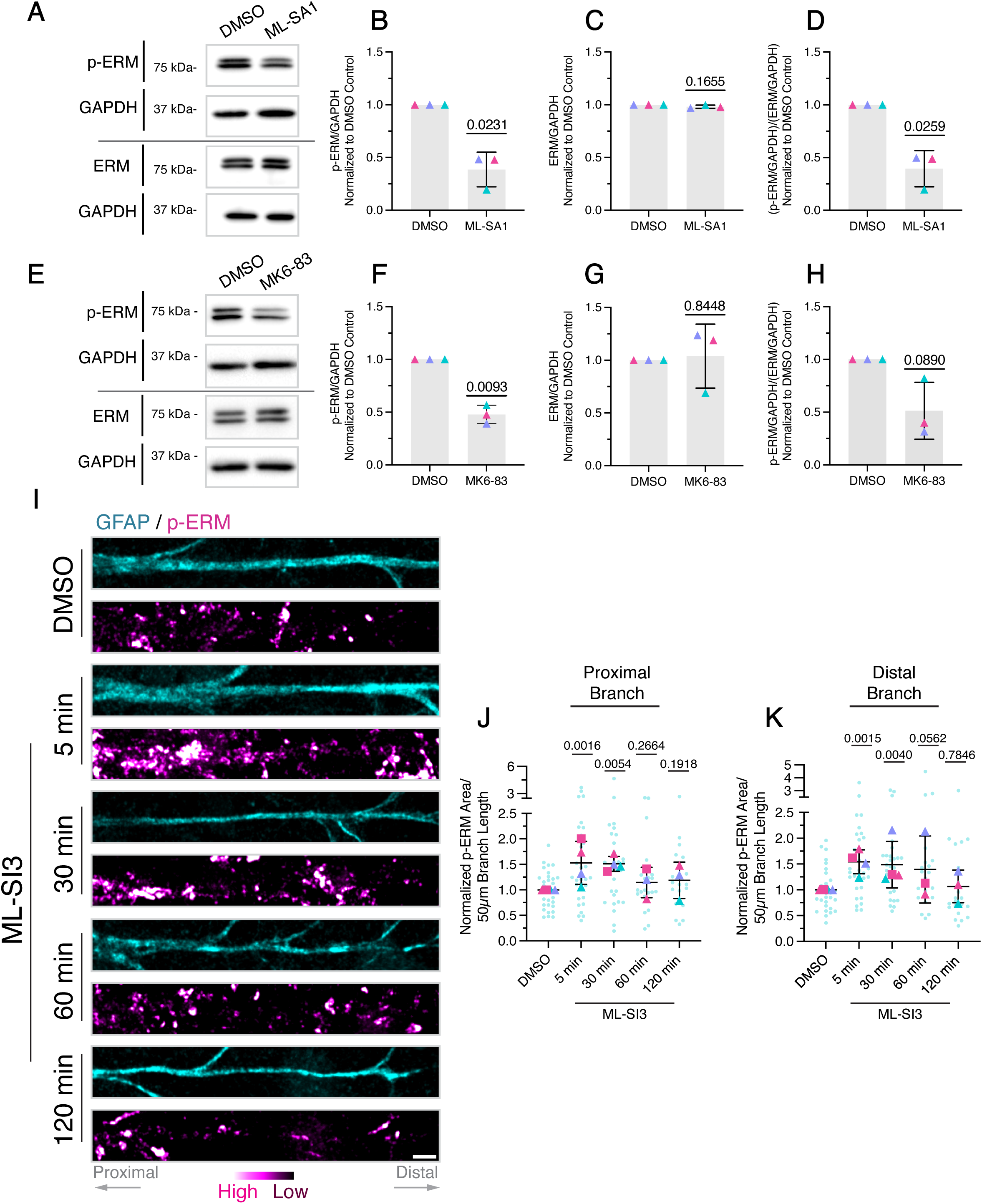
Modulating TRPML1 activity regulates ERM phosphorylation in astrocytes. (A-D) **(A)** Immunoblot analysis of lysates from non-transgenic monocultured astrocytes (DIV7-9) treated with DMSO (solvent control) or ML-SA1 (60 µM) for 5 minutes. Corresponding densitometric analysis of **(B)** p-ERM or **(C)** total ERM levels, normalized to a GAPDH loading control and reported as a fold change relative to the DMSO control. Note p-ERM and ERM proteins appear as doublets by immunoblot. **(D)** Ratio of p-ERM/GAPDH to total ERM/GAPDH, reported as a fold change relative to the DMSO control. Horizontal bars represent the mean of biological replicates ± SD; shown are p-values from a one-sample t-test; N=3 biological replicates. **(E-H)** Immunoblot analysis of lysates from non-transgenic monocultured astrocytes (DIV4-10) treated with DMSO or MK6-83 (30 µM) for 5 minutes. Corresponding densitometric analysis of **(F)** p-ERM or **(G)** total ERM levels, normalized to a GAPDH loading control and reported as a fold change relative to the DMSO control. **(H)** Ratio of p-ERM/GAPDH to total ERM/GAPDH, reported as a fold change relative to the DMSO control. Horizontal bars represent the mean of biological replicates ± SD; shown are p-values from a one-sample t-test; N=3 biological replicates. **(I-K) (I)** Straightened branches from maximum-intensity projections of z-stacks acquired of non-transgenic cocultured astrocytes at DIV7-8; astrocytes were identified by immunostain for GFAP (or S100β, not shown) and counter stained with p-ERM. Cocultures were treated with DMSO (solvent control) for 120 minutes or ML-SI3 (30 µM) for 5, 30, 60, or 120 minutes. p-ERM signal is displayed with identical minimum and maximum intensity settings across treatment conditions. Scale bar, 5 µm. Corresponding quantification of the total area occupied by p-ERM-positive structures within the **(J)** proximal or **(K)** distal 50 µm branch segment, normalized to the DMSO control mean of the corresponding biological replicate. Horizontal bars represent the mean of biological replicates ± SD; shown are p-values from an LME model; N=21-32 branches (one branch per astrocyte) across 3-4 biological replicates.

